# Functional coupling between ribosomal RNA transcription and processing guided by stable transcription factor binding

**DOI:** 10.64898/2026.03.17.712431

**Authors:** Anastasiia Chaban, Nusrat S. Qureshi, Olivier Duss

**Affiliations:** Molecular System Biology Unit, European Molecular Biology Laboratory (EMBL), Heidelberg, Germany; Faculty of Biosciences, Heidelberg University, Heidelberg, Germany

## Abstract

Coordinating ribosomal RNA (rRNA) transcription, folding and processing is essential for bacterial ribosome assembly. Yet, the molecular mechanisms underlying this coordination remain poorly understood. Particularly, whether the rRNA transcription antitermination complex (rrnTAC: NusA, NusG, NusB, NusE, S4, SuhB) orchestrates this coordination remains unclear. Here, we develop a suite of multi-color single-molecule fluorescence microscopy assays to simultaneously visualize, in real-time, rrnTAC assembly dynamics and their effect on transcription and co-transcriptional processing by RNase III. We find that transient (∼1 s) interactions of general transcription factors NusA and NusG with RNAP, and of NusB/E with the rRNA-specific boxBAC element, are stabilized to minutes-long residence by final SuhB recruitment. Stable rrnTAC assembly is required to reduce RNAP pausing and to boost co-transcriptional rRNA processing. Collectively, our results reveal how transcription factor binding dynamics relate to function: transient for messenger RNA transcription and stable for rRNA transcription and processing.

## Introduction

Ribosome assembly is a fundamental process in all cells, consuming the majority of cellular resources under fast growing conditions in *E. coli* ^1^. It is a highly complex, yet efficient, process involving simultaneous transcription of the ribosomal RNA (rRNA), its folding, modification and processing and the binding of ribosomal proteins (r-proteins) to the nascent rRNA to eventually produce a functional ribosome ^2,3^. Despite decades of biochemical and structural investigations, the choreography of these interconnected processes, and whether and how their coupling may boost the efficiency of ribosome assembly remains poorly understood. Indeed, recent attempts to simulate the *in vivo* situation by co-transcribing rRNA in presence of the full complement of ribosomal proteins ^4,5^ or even in presence of cell extract ^6^ show improved efficiencies compared to early reconstitutions executed in absence of rRNA transcription. Nevertheless, *in vitro* ribosome assembly is still an order of magnitudes slower than *in vivo*. Thus, additional mechanisms operate in a living cell and must be recreated in a test tube to quantitatively dissect the molecular mechanism of cooperation between the involved processes.

A critical challenge in bacterial ribosome synthesis is the coordination of three simultaneous processes: rapid transcription of a long primary transcript (pre-rRNA), co-transcriptional folding of the highly structured rRNA, and precise processing by nucleases. The primary processing step that releases the nascent rRNA from the transcription elongation complex is performed by the generally conserved double-stranded RNA (dsRNA)-cleaving endonuclease RNase III ^7^. rRNA processing by RNase III requires the formation of long-range RNA-RNA interactions during transcription elongation, both for releasing the pre-16S and pre-23S rRNA from the single precursor rRNA (1657 nt and 2904 nt of intervening rRNA for pre-16S and pre-23S rRNAs, respectively) ^7–9^. However, understanding how the cells achieve efficient co-transcriptional formation of long-range RNA-RNA interactions is a grand challenge. While intractable *in vivo*, recapitulating the process *in vitro* remains exceedingly difficult as immediate local non-native RNA structure formation upon RNA polymerase (RNAP) exit renders the process fundamentally inefficient ^4,5,10^. Yet, RNase III cleavage *in vivo* occurs within seconds after transcription of the substrate helix ^11^. Thus, it is not clear how the 5’ and 3’ halves of the RNase III substrate helices find each other during active transcription elongation.

To sustain the large amounts of rRNA required for ribosome biogenesis in fast growing cells, transcription of rRNAs is initiated on every operon nearly every second, resulting in high density of transcribing RNAPs on rRNA operons (*rrn*) ^12,13^. A conserved player in rRNA transcription is the ribosomal RNA transcription antitermination complex (rrnTAC) ^14–18^. The rrnTAC is a large protein complex composed of multiple accessory factors, including the Nus proteins (NusA, NusB, NusG, NusE=r-protein S10), homodimeric SuhB, and primary r-protein S4 ^19–21^. Recent cryogenic electron microscopy (cryoEM) structures illustrate how the rrnTAC proteins are bound to RNAP. They also recognize specific elements at the 5’ leader of the rRNA (called N-protein utilization elements or nut elements) which consist of a boxB stem loop followed by boxA and boxC sequences ^22^ (Fig. 1A-C). Functionally, the rrnTAC protects rRNA transcripts from premature transcription termination by the Rho-factor and increases the rates of rRNA synthesis by suppressing RNAP pausing, resulting in an estimated average speed of 90 nucleotides/seconds (nt/s) which is 2-fold faster than transcription of mRNA ^23–25^. However, how the six rrnTAC proteins achieve fast and efficient assembly onto the nascent rRNA is not understood. For example, premature stable incorporation of some of the rrnTAC proteins could lock the nascent rRNA into misfolded conformations, preventing timely assembly of the complete functional complex required for the fast transcription initiation rates of ∼1 event per second ^12,13^. Conversely, if the rrnTAC proteins remain only transiently bound during transcription elongation, they may not effectively shield the nascent RNA from Rho-dependent termination or may not sufficiently reposition NusA to antagonize transcription pausing. Thus, whether transient interactions or stable association of the rrnTAC proteins with RNAP modulate RNAP activity and its interconnected processes is not known. More generally, whether transcription factor interaction dynamics with the transcription machinery differ for rRNA and mRNA remains unclear.

**Figure 1.**
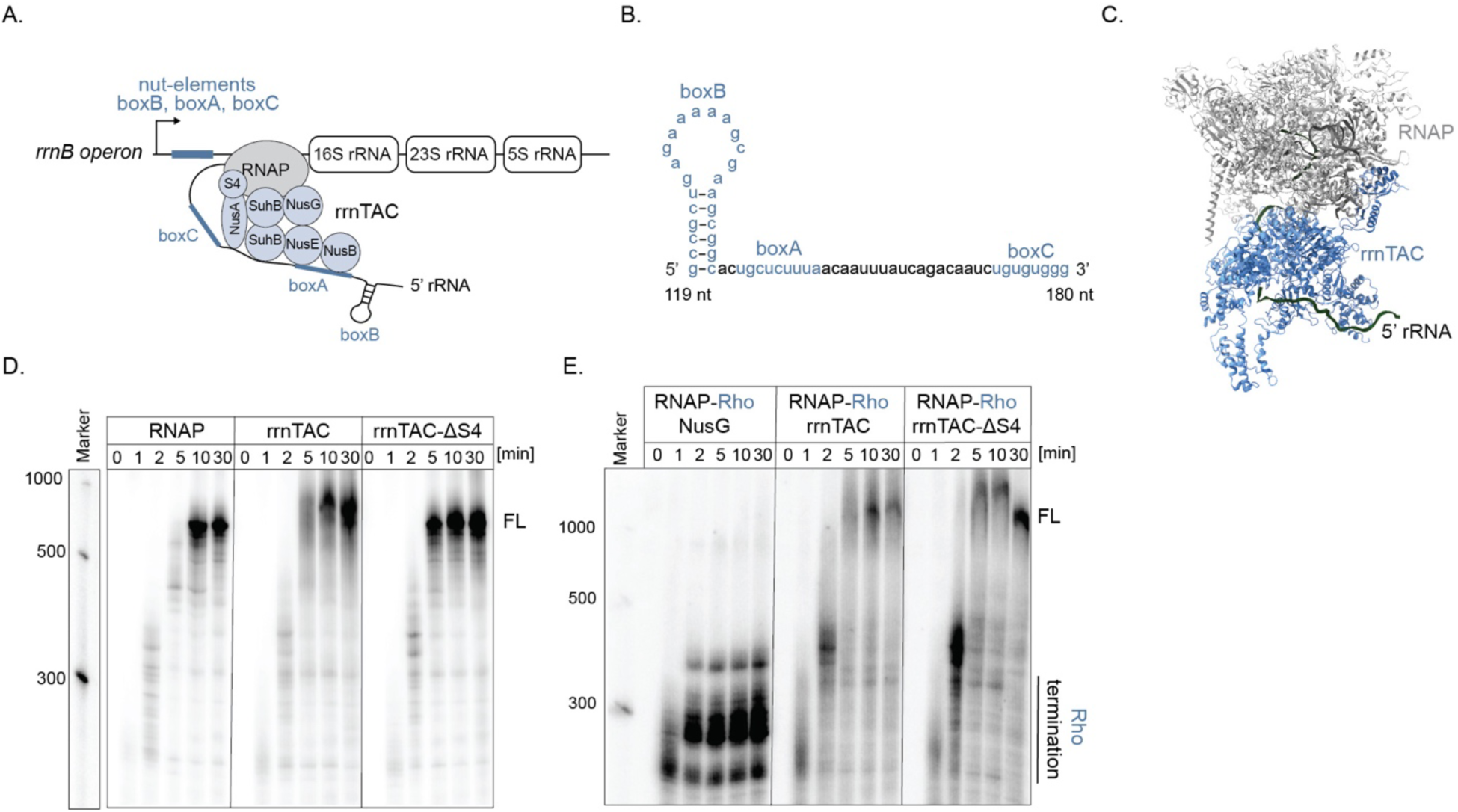
Re-constituted rrnTAC suppresses transcription pausing and Rho transcription termination. (A) Schematic representation of rrnTAC-mediated transcription of pre-16S rRNA. (B) Secondary structure of boxBAC 5’leader rRNA of *rrnB* operon. (C) rrnTAC cryoEM structure (PDB: 6TQO). (D,E) Bulk *in vitro* transcription assays demonstrating rrnTAC activity in promoting transcription elongation (D) and suppressing Rho dependent transcription termination (E). Full gels shown in Fig. S1. Full-length RNA (FL) and Rho termination regions are indicated.

To address these questions, we sought to directly visualize rrnTAC assembly and investigate its potential role in coordinating co-transcriptional rRNA processing. To this end, we developed a suite of multi-color single-molecule fluorescent microscopy assays in which we reconstituted the complete *E. coli* rrnTAC and systematically tracked the real-time binding dynamics of individual rrnTAC components to an active transcription elongation complex (TEC). We show that the general transcription factors NusA and NusG initially transiently associate with RNAP and the NusB-NusE heterodimer transiently binds to boxA at the 5’ end of the RNA. These individually weak protein-RNA and protein-RNAP interactions are rapidly stabilized by the sequential recruitment of SuhB, which transforms a dynamic ensemble of weak interactions into a stable, functional complex within seconds. In contrast, for mRNA transcription, the general transcription factors NusG and NusA only bind transiently, due to the missing boxBAC element. These findings illustrate how the same transcription factors specialize their function by adapting their interaction dynamics with the transcription machinery, being transient (seconds) for mRNA transcription but stable (minutes) for rRNA transcription. Next, by simultaneously monitoring rrnTAC assembly, transcription elongation, and RNase III-mediated rRNA processing of the complete 17S rRNA in a single experiment, we find a drastic increase in co-transcriptional processing efficiency for the subset of active transcription elongation complexes with a completely and stably assembled rrnTAC associated with it. In contrast, inability to form a stable rrnTAC complex results in almost complete loss of co-transcriptional processing. Collectively, these findings provide the first quantitative and mechanistic model of how the abundant and conserved rrnTAC coordinates rRNA transcription, RNA folding, and processing to ensure rapid and robust ribosome assembly.

## Results

### Reconstituted rrnTAC suppresses transcription pausing and Rho transcription termination

To investigate the role of the rrnTAC in transcription and processing of rRNA, we *in vitro* reconstituted the complete *E. coli* rrnTAC from recombinantly purified proteins. To test its activity, we performed single-round bulk *in vitro* transcription assays of the first 900 nucleotides of the pre-rRNA (P1 promoter till end of 5’-domain) (Fig. 1D, S1A-B). Our experiments demonstrate that in presence of all the rrnTAC proteins, yet independent of S4, the ∼900 nucleotides full length transcript is synthesized about twice as fast as in absence of the rrnTAC proteins, in agreement with previous reports ^22^. Next, to test the known activity of the rrnTAC in suppressing Rho-dependent transcription termination ^24^, we added recombinantly expressed Rho and NusG (the latter activates Rho-dependent termination) to our assays and observed strong Rho-dependent transcription termination in absence of the rrnTAC (Fig. 1E, S1B). In presence of the rrnTAC, independent of whether S4 was present, Rho-dependent transcription termination was completely abolished. These data confirm the activity of our reconstituted rrnTAC in promoting transcription elongation and preventing Rho-dependent transcription termination and suggest a minor role for S4 in assisting the known functions of the rrnTAC.

### rrnTAC proteins bind only transiently to isolated TEC and boxBAC element

Having reconstituted the active rrnTAC, we sought to investigate the mechanism of rrnTAC assembly onto the transcription elongation complex (TEC). To simulate the earliest stage during rRNA transcription in which the rrnTAC proteins can start interacting with the transcription machinery, we first tracked the binding of each rrnTAC protein individually to a stalled transcription elongation complex not having yet transcribed the boxBAC RNA element. Notably, this setup also corresponds to the functional transcription elongation complex relevant for transcription of mRNAs which lack the boxBAC sequence. To this end, we prepared a stalled transcription elongation complex (see methods), and immobilized it on a glass surface via the 5’ end of its nascent RNA for single-molecule imaging (Fig. 2A). Informed by the cryoEM structures of the rrnTAC bound TEC (Fig. S2A) ^22^, specific binding of the rrnTAC proteins was detected by FRET between a Cy3 donor dye located on the nascent RNA (13 nt away from RNAP exit channel) and a Cy5 acceptor dye site-specifically incorporated into the rrnTAC protein of interest (see methods). The single-molecule experiments, all performed at 37 °C if not stated otherwise (see methods), were initiated by injecting a Cy5-labeled rrnTAC protein (see specified concentrations in methods) in either absence or presence of all the other unlabeled rrnTAC proteins (Fig. 2B). While NusA and NusG showed multiple transient binding events in absence of the other rrnTAC proteins, little to no binding of the other rrnTAC proteins was detected (Fig. 2C-E). We also determined the NusA and NusG off-rates: k_off_(NusA) = 2.4 +/- 0.2 s^-1^ (bound lifetime = 0.42 +/- 0.04 s); k_off_(NusG) = 0.7 +/- 0.1 s^-1^ (bound lifetime = 1.43 +/- 0.04 s) (Fig. S2C-E). Next, we repeated the experiments by adding all the other rrnTAC proteins to the reaction and found again that only NusA and NusG bound transiently (k_off_(NusA)= 1.4 +/- 0.1 s^-1^ (bound lifetime = 0.71 +/- 0.05 s); k_off_(NusG)= 0.67 +/- 0.01 s^-1^ (bound lifetime = 1.49 +/- 0.02 s)), while we could not detect binding for the other rrnTAC proteins (only events detected with a bound dwell of > 100 ms) (Fig. 2C-E, S2C-E).

**Figure 2.**
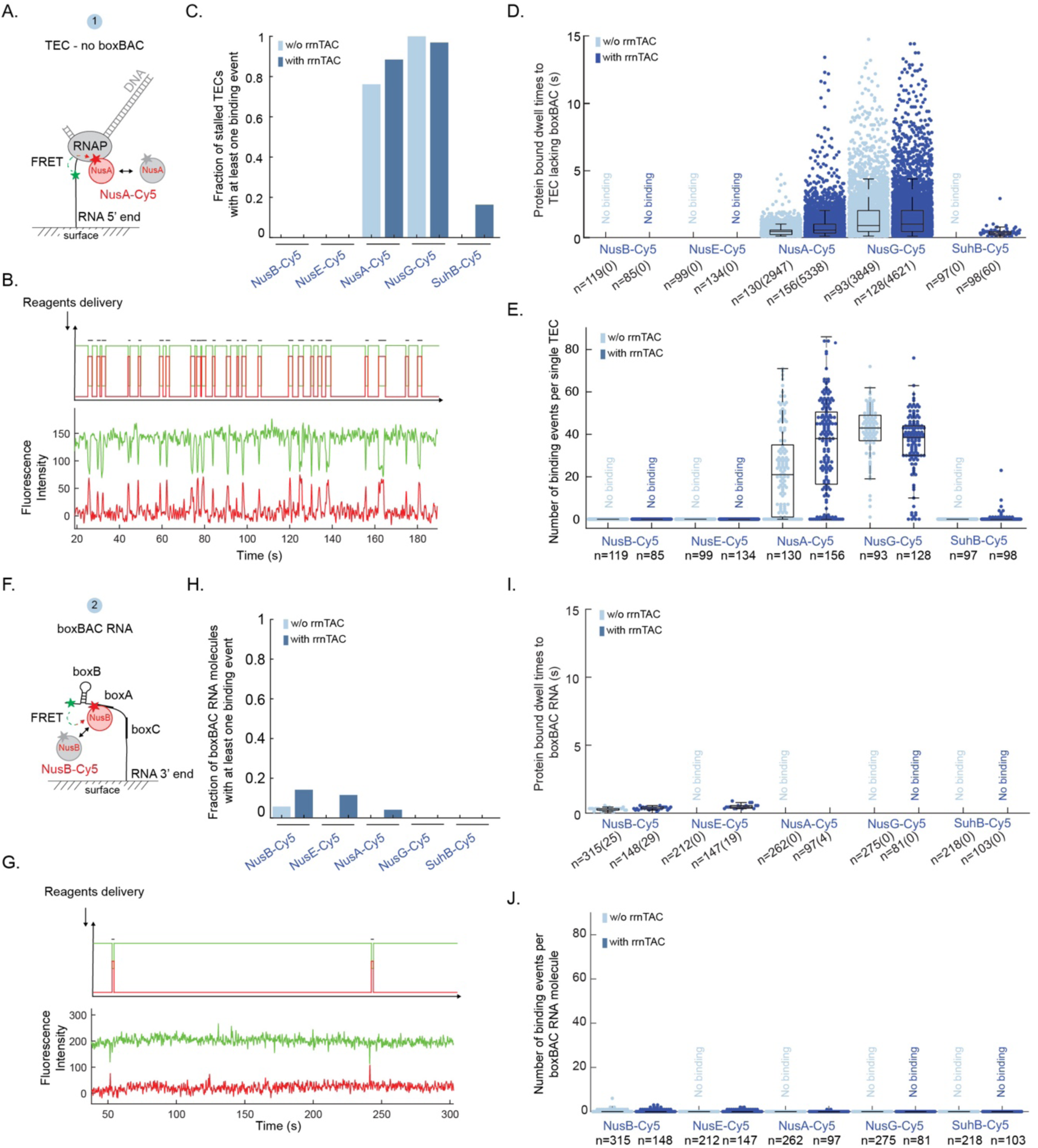
rrnTAC proteins bind only transiently to isolated TEC or boxBAC element. (A) Experimental setup to measure binding of rrnTAC proteins to the stalled TEC lacking the boxBAC RNA element. (B) Upper panel: schematic representation of a single-molecule trace. Bottom panel: representative single-molecule trace of NusA-Cy5 binding to the stalled Cy3-TEC. (C) Fraction of the stalled TECs with at least one rrnTAC protein binding event. (D,E) Beeswarm plot with overlaid boxplot representing all protein-bound dwell times of < 15 seconds (D) or the number of binding events per trace (E) for the different Cy5-labeled rrnTAC proteins binding to the TEC lacking the boxBAC RNA element. (F) Experimental setup to measure binding of rrnTAC proteins to RNA encoding the boxBAC element. (G) Upper panel: schematic representation of a single-molecule trace. Bottom panel: representative single-molecule trace of NusB-Cy5 binding to the Cy3-boxBAC RNA. (H) Fraction of boxBAC RNA molecules with at least one rrnTAC protein binding event. (I,J) Beeswarm plot with overlaid boxplot representing all protein-bound dwell times of < 15 seconds (I) or the number of binding events per trace (J) for the different Cy5-labeled rrnTAC proteins binding to the boxBAC RNA. Number of molecules (n) and events (in brackets) is indicated.

Next, we investigated which rrnTAC proteins can interact with the boxBAC RNA independently from RNAP, to simulate the situation in which the rrnTAC proteins could seed the assembly of the rrnTAC on the boxBAC element once it has emerged from the RNAP. To this end, we immobilized a pre-transcribed RNA encoding the boxBAC element, labeled with a Cy3 donor dye, to the glass surface for single-molecule imaging and added again combinations of Cy5-acceptor labeled rrnTAC proteins in presence or absence of the other unlabeled rrnTAC proteins (Fig. 2F). We hardly detected binding for any of the proteins, except for very rare and transient binding events for NusE and NusB at 37 °C (Fig. 2F-J, S2F). Repeating our experiments at 21 °C (Fig. S2G-J), the binding became more frequent and we could determine off-rates of k_off_(NusB) = 1.5 +/- 0.1 s^-1^, k_off_(NusE) = 1.3 +/- 0.1 s^-1^ (Fig. S2K), in agreement with previously reported interactions of NusE/B with boxBAC RNA ^26^. Altogether, our data demonstrate that only NusA and NusG transiently bind to the transcription elongation complex lacking the boxBAC RNA element. In contrast, only NusB and NusE very weakly bind the isolated boxBAC RNA. These data are consistent with a model in which NusA and NusG seed assembly on the RNAP while NusE and NusB seed assembly on the boxBAC RNA. Furthermore, these findings demonstrate that the transcription factors NusA and NusG bind only transiently to an mRNA transcription elongation complex.

### Both TEC and boxBAC elements are required for stable rrnTAC formation

Having shown that none of the rrnTAC proteins bind stably to either the RNAP or boxBAC RNA element alone, we next investigated the binding dynamics of the rrnTAC proteins to a transcription elongation complex in which the boxBAC element has fully emerged from the RNAP exit tunnel. To this end, we prepared a stalled transcription elongation complex containing the boxBAC RNA using an artificially assembled RNA:DNA scaffold bound by RNAP (Fig. 3A, Fig. S3A). The RNA in the scaffold encoded the entire boxBAC leader and was labeled with a Cy3-donor dye to detect binding of the Cy5-labeled rrnTAC proteins to the transcription elongation complex via FRET as in the previous assays. The DNA in the scaffold encoded the 5’ domain of 16S rRNA of the native *rrnB* operon sequence, followed by a single transcription terminator and 2xCy3.5 dyes to track transcription. This setup allowed us to verify that each artificially assembled TEC scaffold included in our analysis was functional, by following a 2-step procedure (Fig. 3A): first, we monitored rrnTAC protein binding by injecting individual Cy5-labeled rrnTAC proteins either in absence or presence of all the other unlabeled rrnTAC protein factors. Second, to select only the transcriptionally active molecules for data evaluation, we injected NTPs and followed transcription elongation by detecting a pattern of an initial Cy3.5 signal intensity increase (due to the approach of the Cy3.5 dyes to the surface during transcription elongation) followed by loss of the Cy3.5 signal (due to DNA template dissociation at the transcription terminator) ^27^.

**Figure 3.**
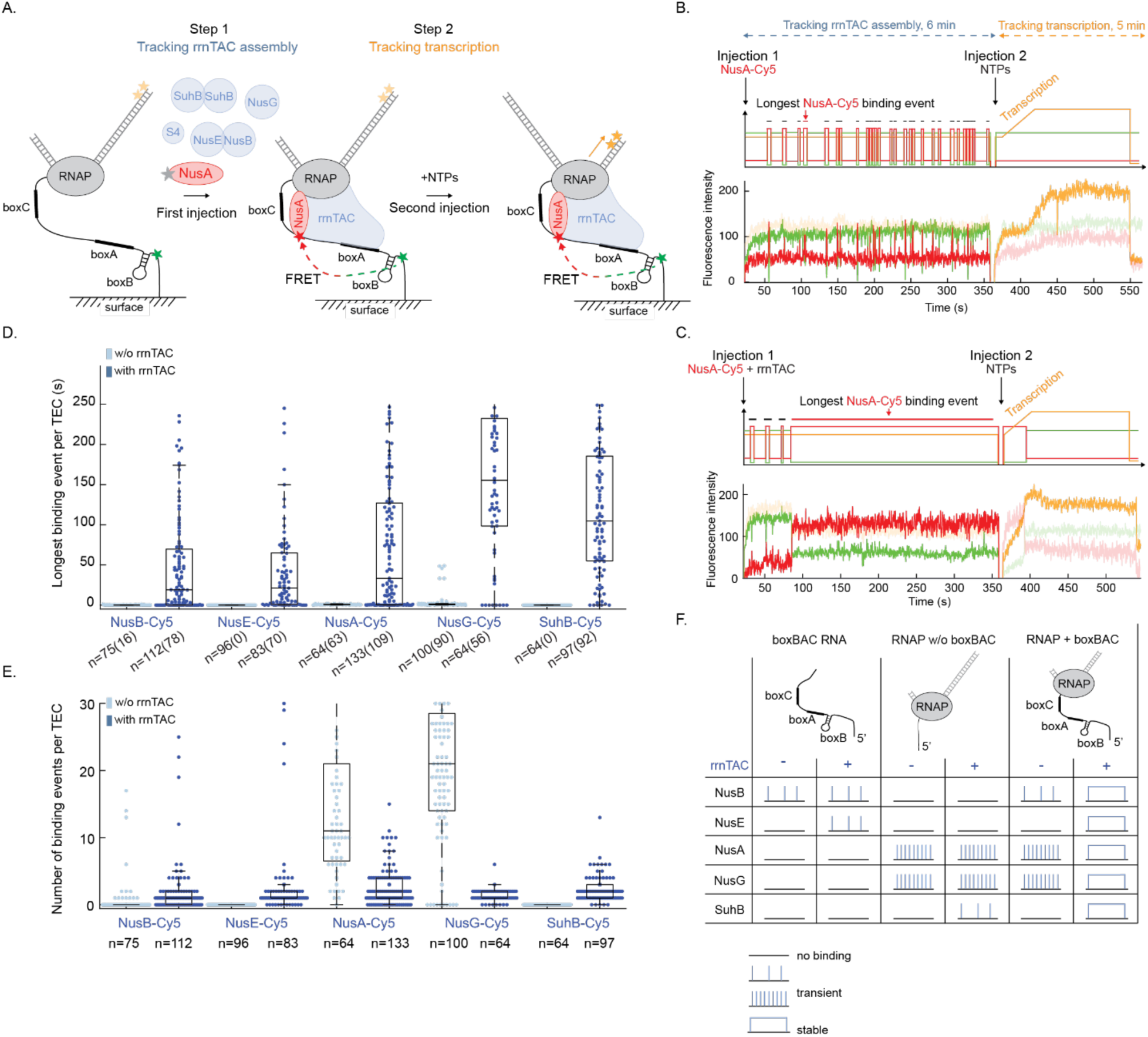
Both TEC and boxBAC element are required for stable rrnTAC formation. (A) Experimental setup to study rrnTAC assembly on the boxBAC-containing TEC. (B,C) Schematic (top) and representative experimental (bottom) single-molecule trace. Example: 10 nM NusA-Cy5 in absence (B) or presence of 400 nM of the other rrnTAC proteins (C). (D,E) Beeswarm plot with overlaid boxplot representing the longest binding event per trace (D) or the number of binding events per trace (E) for the different Cy5-labeled rrnTAC proteins binding to the TEC containing the boxBAC RNA element. Only data of <250 s (D) or <31 events (E) are shown. Longest binding event for molecules that did not show any binding are plotted as zero. Number of molecules (n) and events (in brackets) are indicated. (F) Overview of rrnTAC protein binding kinetics to various constructs.

In presence of the boxBAC-containing TEC, we again found that only NusA and NusG, and very weakly NusB, bind transiently in absence of the other rrnTAC proteins (Fig. 3B, Fig. S3B). However, upon addition of all the other rrnTAC proteins, we observed a drastic increase in bound-dwell times for all the rrnTAC proteins. The traces initially showed none or few transient events followed by a minutes-long event, often limited by photobleaching (Fig. 3C Fig. S3B). We interpret the initial transient binding as unstable rrnTAC assembly intermediates and the stable event as the fully assembled rrnTAC. To quantify the binding dynamics across the different experimental conditions and enriching for the assembled state, we plotted the longest binding event (Fig. 3D) and the total number of binding events for each single-molecule trace (Fig. 3E). This demonstrates orders of magnitude longer bound dwell times of the rrnTAC proteins in presence of the other rrnTAC proteins compared to their isolated binding. Altogether, our data show that the rrnTAC can only stably assemble in the presence of both RNAP and the rRNA-specific boxBAC RNA element and thus, illustrate that the general transcription factors NusG and NusA engage in transient interactions with mRNA transcription elongation complexes (lacking the boxBAC element) but stably associate during rRNA transcription, mediated by at least a subset of the other rrnTAC proteins (Fig. 3F).

### SuhB incorporation requires prior binding of NusA, NusG and the NusB-NusE heterodimer

For SuhB, we did not detect any binding to the boxBAC containing TEC in absence of the other rrnTAC proteins. Yet, functional transcription assays supported the importance of SuhB: absence of SuhB failed suppressing transcription pausing, in contrast to the situation when all rrnTAC proteins are present (Fig. S1B, middle gel). Therefore, we focused next on understanding the role of the SuhB homodimer in the rrnTAC assembly. After having verified that SuhB is present as a homodimer at concentrations below our experimental conditions (Fig. S2B), in agreement with structural data ^24^, we first quantified which rrnTAC proteins are required for SuhB incorporation into the rrnTAC. To this end, we measured the real-time binding dynamics of SuhB-Cy5 to the boxBAC containing TEC and systematically omitted one rrnTAC protein at a time. Omitting NusA completely abolished SuhB incorporation, highlighting the absolute requirement for NusA (Fig. 4A). Similarly, omitting NusG prevented SuhB recruitment, consistent with molecular interactions between NusG and SuhB in the rrnTAC/TEC structure ^24^. In contrast, omitting r-protein S4 did not affect SuhB recruitment efficiency, bound dwell times and recruitment speed (Fig. S5A-C).

**Figure 4.**
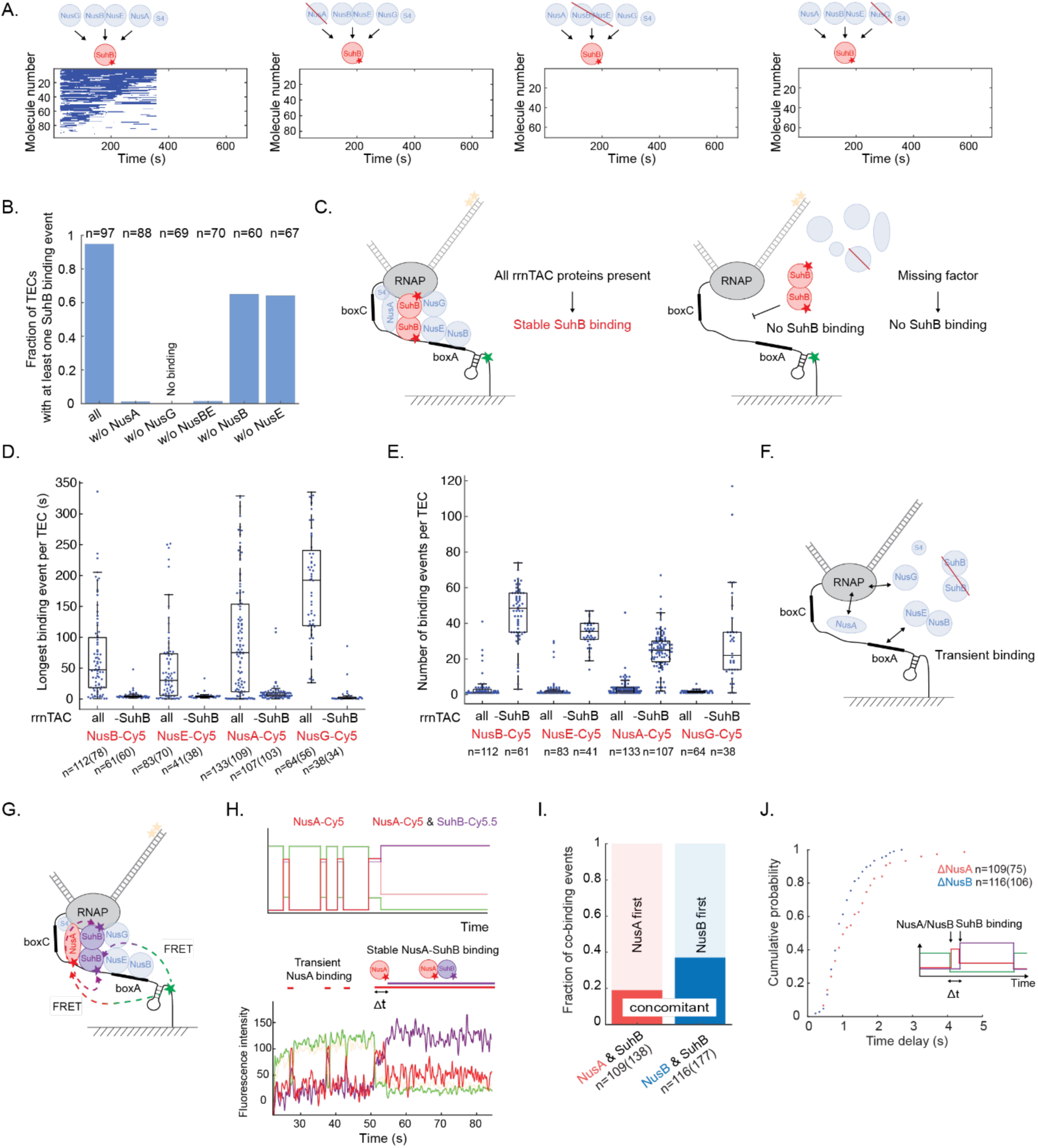
SuhB incorporates in rrnTAC last, stabilizing transient binding of NusA, NusG and NusB-NusE heterodimer. (A) Rastergrams with each row representing a single stalled TEC to which SuhB binding events are represented as blue bars, in presence of various unlabeled rrnTAC proteins. (B) Fraction of TECs with at least one SuhB-Cy5 binding event in presence of various unlabeled rrnTAC proteins as indicated. (C) Schematic illustrating that SuhB only stably incorporates in presence of all other rrnTAC proteins. (D,E) Beeswarm plot with overlaid boxplot representing the longest binding event per trace (D) or the number of binding events per trace (E) for the different Cy5-labeled rrnTAC proteins binding to the boxBAC-containing TEC in presence of all rrnTAC proteins but either in presence or absence of SuhB. Longest binding event for molecules that did not show any binding are plotted as zero. (F) Schematic illustrating the transient binding of all rrnTAC proteins in absence of SuhB. (G-J) Simultaneous tracking of two rrnTAC proteins (NusA/B-Cy5 and SuhB-Cy5.5) allows quantification of order and kinetics of assembly: experimental setup (G); schematic (top) and representative experimental (bottom) single-molecule trace (H); quantification of order of binding (I). Events too fast to be detected sequentially were assigned as “concomitant”; cumulative probability showing the time delay between NusA/NusB and SuhB binding (J). Number of molecules (n) and events (in brackets) is indicated for all experiments.

To test the role of the boxA-binding NusE/B heterodimer ^28^, for which we confirmed a heterodimeric state (Fig. S4), we omitted NusB and NusE simultaneously. We also detected complete loss of SuhB binding under these conditions. However, omitting either NusB or NusE individually resulted in stable SuhB incorporation (Fig. S5A-B) but in a reduced fraction of complexes (Fig. 4B). Furthermore, SuhB recruitment kinetics were dramatically reduced: the median SuhB arrival time increased to 110 s (no NusB) or 101 s (no NusE), compared to 5 s in presence of all the rrnTAC proteins (Fig. S5C). These data are in line with *in vivo* observations, where deletion of *nusB* impairs rRNA synthesis, yet defects that can be rescued by *nusE* overexpression ^19,29–31^. The *in vivo* phenotypes are mechanistically supported by our demonstration that loss of NusB results in slower and less efficient SuhB incorporation, thereby drastically delaying rrnTAC maturation. Overall, these experiments demonstrate that all rrnTAC proteins, except of S4, are required for efficient and functional SuhB recruitment to the boxBAC containing TEC (Fig. 4C).

### All proteins are required for assembly of a stable and functional rrnTAC

Having established that all rrnTAC proteins are required for SuhB recruitment to the rrnTAC, we wondered whether the rrnTAC can also stably assemble in the absence of SuhB. Upon omitting SuhB, all other rrnTAC proteins bound the boxBAC containing stalled transcription complex only transiently (shorter dwell times, increased numbers of events per molecule; Fig. 4D-F). To test whether the requirement for all rrnTAC proteins is general for stable rrnTAC complex formation, we used NusA binding as another reporter for complex stability. Sequential omission of each of the other rrnTAC proteins revealed that stable NusA incorporation, and thus stable assembly of the full rrnTAC, only occurs when the full complement of the rrnTAC proteins is present (Fig. S5D). Overall, our data underscore the absolute requirement for all rrnTAC proteins but S4, for stabilizing the complex, converting dynamic protein recruitment into stable and functional incorporation of the rrnTAC components.

### Two-component tracking reveals rapid rrnTAC assembly kinetics

We next quantified the kinetics of rrnTAC assembly by tracking one protein initiating assembly on the RNAP (NusA) or the boxBAC RNA element (NusB) together with SuhB, which completes assembly. To this end, we first recorded simultaneous binding of differently labeled NusA and SuhB proteins to the stalled elongation complex in real-time (Fig. 4G-H, Fig. S6A). At 10 nM NusA-Cy5 and 50 nM SuhB-Cy5.5 in presence of the other unlabeled rrnTAC proteins, we quantified the order of protein binding and their kinetics. The majority of the co-binding events (81%) showed NusA binding first followed by SuhB binding with a median delay of 1 s (Fig. 4I-J). The rest of co-binding events either had too fast sequential binding to allow resolving them with our experiments (<100 ms delay) or occurred as binding of heterodimers as suggested by direct interactions between NusA and SuhB ^24^. Similarly, quantifying the relative binding kinetics of the NusB/E heterodimer and SuhB (Fig. S6B) using 50 nM NusB-Cy5 and 50 nM SuhB-Cy5.5 (in presence of the other unlabeled rrnTAC proteins), we found NusB/E binding first in 63 % of the events (Fig. 4I), followed by a median delay of 1 s before SuhB incorporation, with the remaining co-binding events occurring too fast to detect the sequential binding (Fig. 4J). Overall, our data are compatible with a model in which transient binding of NusA and NusG to the RNAP and of NusB/E to the boxBAC rRNA element is subsequently stabilized by SuhB to complete rrnTAC assembly within about a second.

### Stably assembled rrnTAC increases transcription speed two-fold

Having established that all the rrnTAC proteins, except of S4, are required for complete and stable rrnTAC assembly, we wondered whether stable rrnTAC assembly is a determinant for rrnTAC-mediated increase in transcription speed. We therefore quantified the transcription elongation rates for completely versus incompletely assembled rrnTACs. To this end, we immobilized a TEC scaffold (with rrnTAC proteins +/-SuhB) containing the nascent boxBAC sequence, site-specifically labeled Cy5-RNAP and a DNA template coding for the entire 16S rRNA, followed by a transcription terminator and two Cy3.5 dyes (Fig. S7A). This labeling strategy allowed us to identify the timepoint at which the RNAP-Cy5 has completed transcription of the full 16S rRNA by appearance of a Cy3.5-Cy5 FRET (Fig. S7B), thus providing the transcription time for each single TEC. We observed an approximate two-fold increase in transcription rate for TECs containing a pre-assembled rrnTAC (median 171 nt/s at 1 mM NTPs) compared to TECs only containing transient rrnTAC assemblies, observed when omitting SuhB (99 nt/s) (Fig. S7C). This single-molecule result explains the bulk transcription assays (Fig. 1D, S1B), collectively revealing that stable rrnTAC assembly via SuhB is required to increase the transcription elongation rate.

### RNase III requires native helix for productive cleavage

Having investigated the mechanism of complete assembly of the rrnTAC onto the boxBAC containing TEC, we sought to investigate how the rrnTAC might affect co-transcriptional processing by RNase III. As a first step to study these potentially interconnected processes, we developed a simplified assay to track RNase III binding and subsequent cleavage in real-time with a minimal pre-transcribed target RNA. We cloned and purified RNase III with an N-terminal ybbR tag for subsequent fluorescent labeling (Fig. 5A). RNase III acts as a homodimer and cleaves double-stranded RNA (dsRNA) helices of minimal length of 22 nt leaving a 2-nt overhang at the 3’- end of the products ^32,33^. Thus, as a substrate for cleavage, we *in vitro* transcribed an RNA encoding the native RNase III substrate helix of pre-16S rRNA. Bulk RNA cleavage assays of this substrate showed efficient cleavage, verifying activity of our tagged RNase III (Fig. 5B, S8A).

**Figure 5.**
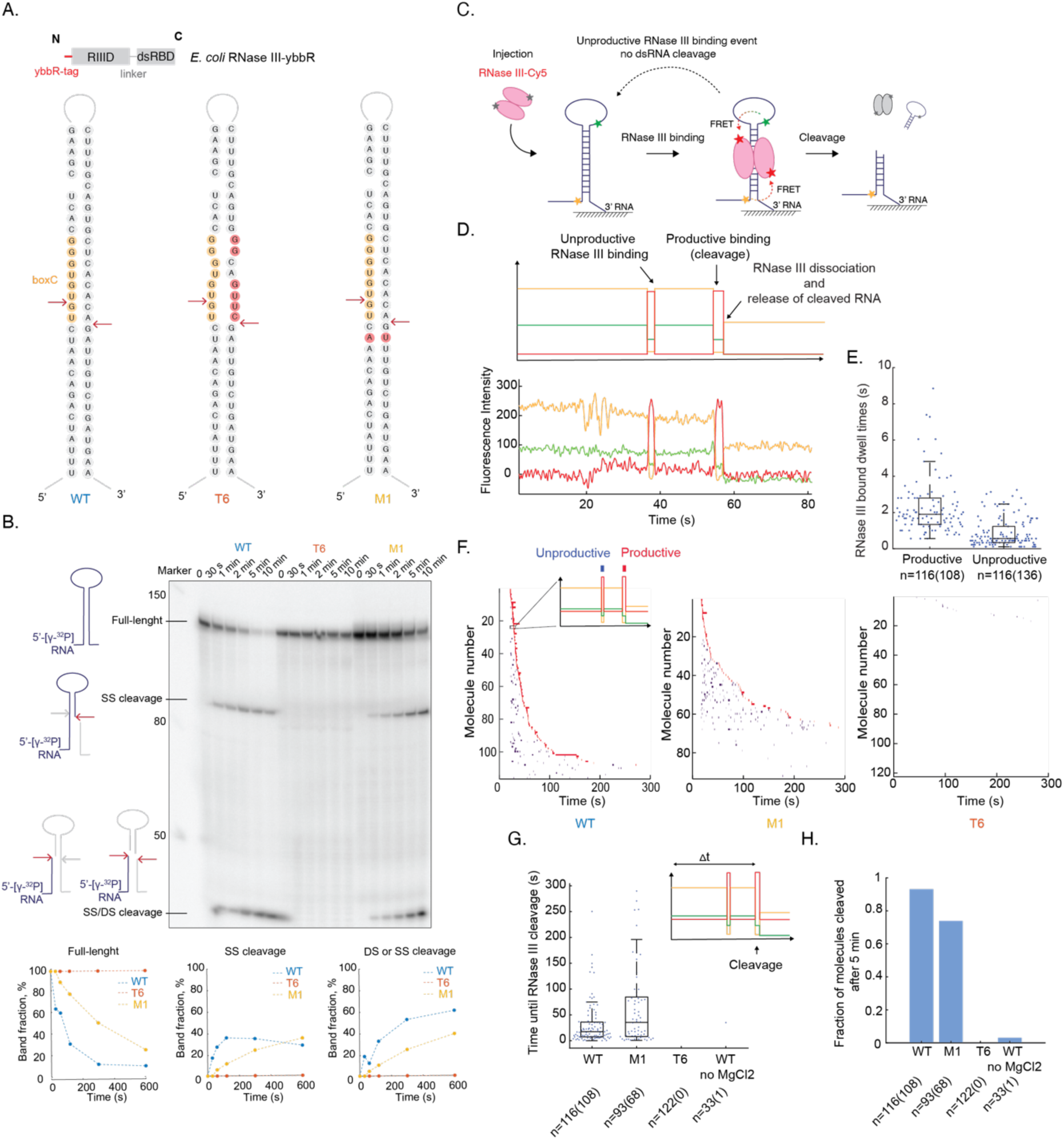
RNase III requires native helix for efficient cleavage. (A) Construct overview. (B) Bulk RNase III cleavage assay demonstrating single-site (SS) and double-site (DS) cleavage. Reactions performed at 3.5 mM MgCl_2_ and 21 °C. (C-E) Single-molecule assay for detection of RNase III binding and scoring for productive versus unproductive cleavage: experimental setup (C); schematic (top) and representative experimental (bottom) single-molecule trace (D); beeswarm plot and overlaid boxplot for RNase III-bound dwell times for productive versus unproductive binding (E). (F-H) Quantification of RNase III cleavage for different mutants: rastergrams with each row representing a single RNA molecule with unproductive (blue) and productive (red) RNase III binding events shown as colored bars (F); beeswarm plot and overlaid boxplot of time until productive binding event (G); bar plot representing fraction of molecules cleaved after 5 minutes. Conditions for single-molecule assays: 20 nM RNase III-Cy5, 3.5 mM MgCl_2_, 21 °C. Number of molecules (n) and events (in brackets) is indicated for all experiments.

Next, to simultaneously track RNase III binding to and cleavage of single substrate RNA molecules in real-time, we immobilized the pre-transcribed and pre-folded model RNA to a glass surface via its 3’ end for single-molecule imaging (Fig. 5C). We labeled the substrate RNA with two donor dyes, a Cy3.5 dye at its 5’end and a Cy3 dye in the internal RNA loop. This labeling strategy allowed us to detect specific binding of accepter-labeled RNase III-Cy5 via FRET to either of the two donor dyes and in addition, to detect whether the respective RNase III binding event lead to productive cleavage, in which case the Cy3 dye in the loop would disappear upon successful cleavage (Fig. 5C). To initiate the reaction, we added 20 nM RNase III-Cy5 to the immobilized target RNA. Under these conditions, our single-molecule traces (Fig. 5D) showed two types of RNase III binding events: about 50 % of the events (Fig. 5E) showed RNase III binding followed by immediate Cy3-dye disappearance upon RNase III dissociation, which we assign as productive RNase III binding events. The other 50% of the RNase III binding events did not show Cy3-dye disappearance, events which we assign as unproductive binding events (detailed below). The median RNase III-bound dwell times were about 3-fold longer for productive (median = 1.9 s) versus unproductive (median = 0.6 s) binding events (Fig. 5E).

In order to determine the molecular basis for the unproductive binding events, we performed bulk cleavage assays. A substantial fraction of RNA molecules showed only a single-site cleavage at earlier time points (Fig. 5B), a fraction which decreased over time or by increasing the temperature (Fig 5B; S8A). We therefore conclude that at least a fraction of the “non-productive” RNase III cleavage events results from RNase III molecules which cleave only at one site of the dsRNA and dissociate prematurely from the target RNA before completing the second cleavage. In support of this, 25 % of the RNA molecules show at least one unproductive binding event before the successful cleavage event under these conditions (Fig. S8C).

Having in hand an assay to separately quantify the number of unproductive binding events until cleavage and to determine the time required until cleavage for each RNA molecule, we next investigated how mutations affect the cleavage dynamics. To this end, we generated two mutant RNAs: the M1 RNA mutant encoded a native helix with a single swapped pair of nucleotides and the T6 RNA mutant encoded six base-pair mismatches that partially disrupted the native helix near the cleavage site and completely abolishes the formation of active ribosomes *in vivo* (Warner et al., 2023). Compared to WT, the M1 RNA mutant showed an approximately 2-fold longer delay until the appearance of a productive cleavage event (median delay till cleavage for WT: 17 s; M1 mutant: 35 s), leading to a smaller fraction of the molecules being processed during the experimental time (Fig. 5F-H). Interestingly, the time required for RNase III to bind the substrate was similar to the WT RNA, but slower processing occurred due to an increased number of unproductive binding events prior the cleavage compared to the WT RNA (Fig. 5G, Fig. S8B-C). For the T6 RNA mutant, we observed only few unproductive binding events and no cleavage, indicating that even slightly disrupting the substrate helix drastically reduces RNase III binding and completely abolishes cleavage. This is in agreement with the bulk cleavage assay showing slower processing of the M1 RNA mutant and the complete lack of processing for the T6 RNA mutant (Fig. 5B, Fig. S8A). Overall, our data demonstrate that RNase III processing of the RNA can proceed through unproductive binding events, some of which lead to cleavage at only one of the two strands, while other events perform both cleavage steps during a single RNase III binding event. Furthermore, RNase III binding and cleavage is abolished if the substrate helix is not correctly formed, demonstrating that RNase III binding can be used as a reporter for native RNase III substrate helix formation.

### Assembled rrnTAC boosts co-transcriptional RNase III processing efficiency

Having established two separate single-molecule assays to either track the complete assembly of the rrnTAC (Fig. 3 and Fig. 4) or to detect RNase III binding and processing in real-time (Fig. 5), we next aimed at combining the two systems into one in order to investigate and directly image how rrnTAC assembly dynamics and composition may affect co-transcriptional rRNA processing by RNase III (Fig. 6). More specifically, we asked the questions: does the rrnTAC increase rRNA processing efficiency by RNase III? If so, is the complete and stably assembled rrnTAC required or could a transient assembly lacking one of the factors perform the same function? In order to address these questions, we followed a two-step approach (Fig. 6C): step 1 – NusA-Cy5.5 in presence of all the other rrnTAC proteins was added to an immobilized boxBAC-containing transcription elongation scaffold in order to pre-assemble the rrnTAC. Stable assembly of the rrnTAC was identified at the single-molecule level by the bound dwell time of NusA-Cy5.5 since NusA (or any other rrnTAC protein) binds stably only when NusG, SuhB and the NusB/NusE heterodimer are present (Fig. 6D,E, S5D, Fig. 4). Step 2 – transcription elongation of the rRNA and subsequent co-transcriptional RNase III processing was initiated by addition of RNase III-Cy5 and specified concentrations of NTPs. To detect specific binding of acceptor-labeled NusA-Cy5.5 (step 1) and RNase III-Cy5 (step 2) to the transcription elongation complex, we constructed and assembled a stalled transcription elongation scaffold containing a nascent RNA consisting of the complete boxBAC sequence, followed by the 5’ half of the RNase III cleavage helix and a site-specifically introduced Cy3-donor dye (introduced by multiple segmental labeling of RNA; see methods) just upstream the RNAP exit tunnel (Fig. 6A,C). The scaffold DNA coded for the entire 5’domain of the 16S rRNA followed by the 3’-half of the RNase III helix and ending with a spacer, terminator and two Cy3.5 dyes to monitor transcription elongation in real-time (Fig. 6B-C; see methods for details).

**Figure 6.**
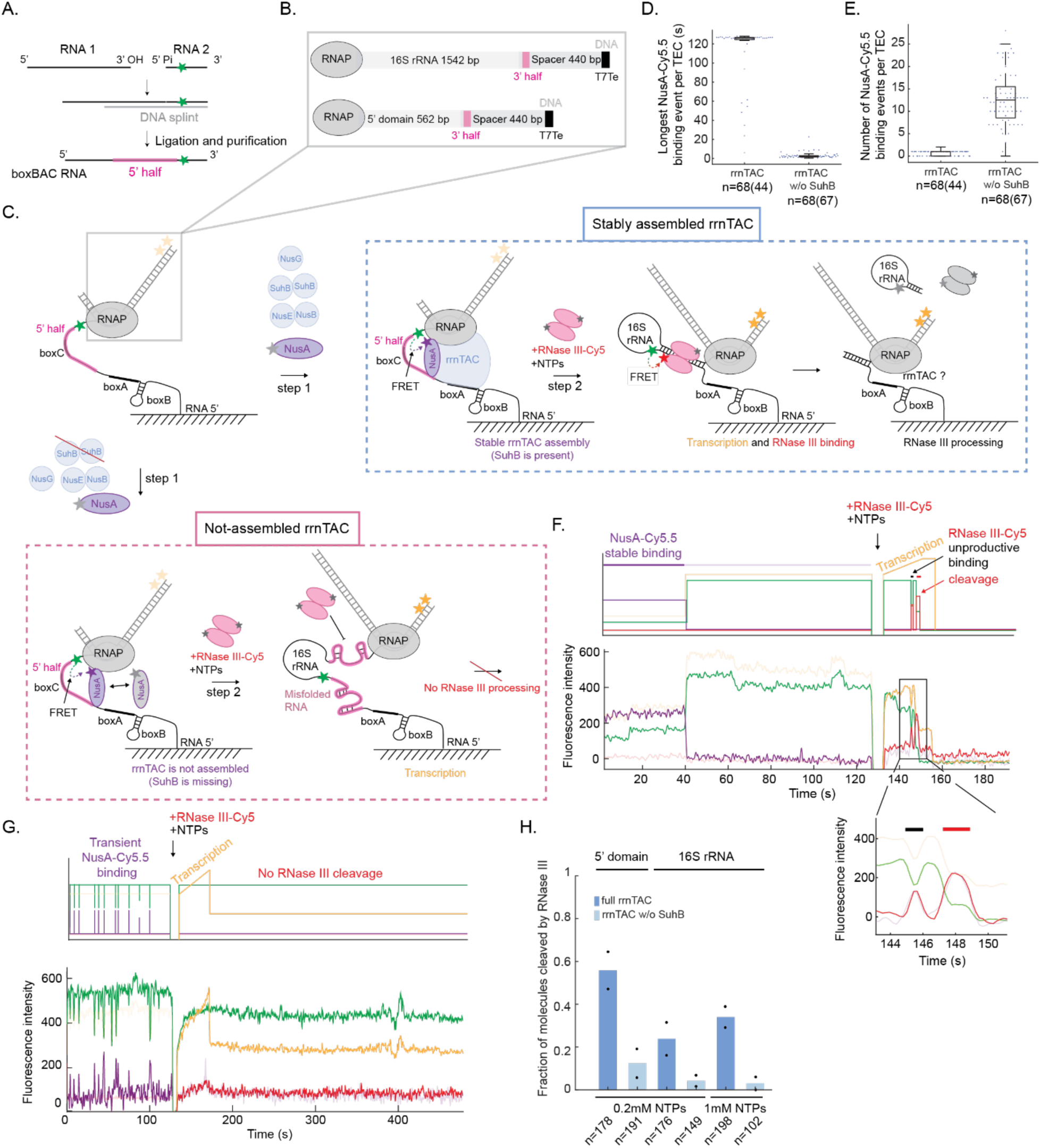
Assembled rrnTAC boosts co-transcriptional RNase III processing efficiency. (A-C) Experimental design: generation of nascent RNA by splinted ligation (A); DNA template design (B); single-molecule experiment to simultaneously detect rrnTAC assembly (step 1) and co-transcriptional RNase III cleavage (step 2) (C). (D,E) Beeswarm plot and overlaid boxplot of the longest NusA-Cy5.5 binding event (D) and number of binding events per trace (E) for assembled and not-assembled rrnTAC (lacking SuhB). For assembled rrnTAC, the bound-dwells are limited by the second injection (<125 s). (F,G) Schematic (top) and representative experimental (bottom) single-molecule trace for assembled (F) or non-assembled (lacking SuhB) rrnTAC (G). (H) Fraction of nascent RNA molecules cleaved by RNase III. Black dots represent values from two biological replicates. Number of molecules (n) and events (in brackets) is indicated for all experiments.

In presence of all rrnTAC proteins (Fig. 6C blue dotted box), ∼75% of the TECs with active transcription (Fig. S9A) showed a single stable NusA-Cy5.5 binding event (Fig. 6D-F, S9B), which we assign to TECs with completely assembled rrnTAC complexes. The remaining molecules with active transcription did not show NusA binding at all. We suggest that the majority of these represent fully assembled rrnTAC complexes bound to an unlabeled NusA molecule. In strong contrast, when omitting SuhB from the reaction (Fig. 6C pink dotted box), almost all the transcriptionally active TECs showed multiple transient NusA-Cy5.5 binding events (Fig. 6D-E,G and S9A,C). We assign these TECs as not having a fully assembled rrnTAC (Fig. 4D-E).

Next, we quantified which fraction of the transcribing TECs lead to successful RNase III processing either when the rrnTAC was completely and stably assembled (Fig. 6C, 6F) or when the rrnTAC was incomplete and transient (Fig. 6C, 6G). Productive RNase III binding events showed simultaneous loss of RNase III - Cy5 and RNA-Cy3 signal, indicating release of the nascent RNA (Fig. 6C,6F, S9B). At 0.2 mM NTPs and transcription of the 5’ domain of the 16S rRNA, 56 % of the nascent rRNAs were processed by RNase III, while the fraction dropped almost 5-fold to 12 % for incompletely TEC-associated rrnTACs (Fig. 6H). These results demonstrate a strong effect of the stably assembled rrnTAC on co-transcriptional processing by RNase III, but also indicate that folding of the long-range helix still occurred also in absence of a completely assembled rrnTAC. We further confirmed the specificity of RNase III cleavage - acting only on its natural target site - by performing bulk *in vitro* transcription assays of the same construct under the same conditions (Fig. S10).

We repeated our experiments at physiological NTP concentrations and with the full native pre-17S rRNA, representing the complete natural substrate for co-transcriptional processing. In presence of the completely assembled rrnTAC, not affected by S4 (Fig. S9D), RNase III processing decreased to about ∼34 %, in agreement with co-transcriptional folding becoming more challenging with increasing RNA length (Duss et al., 2019). However, in absence of the fully assembled rrnTAC, RNase III processing was almost completely abolished (Fig. 6H). This dramatic increase in co-transcriptional processing in the presence of the stable and completely assembled rrnTAC illustrates that the complete and stable rrnTAC is key for co-transcriptional chaperoning of the long-range RNase III substrate helix, thus boosting co-transcriptional rRNA processing efficiency.

## Discussion

### Model for rrnTAC-mediated co-transcriptional rRNA processing

Ribosome assembly has been intensively studied over the past decades. However, still little is known about the kinetics of early rRNA transcription, folding and processing, and the mechanisms of functional interconnectivity between these processes. Here, we *in vitro* reconstitute and describe early ribosome synthesis using single-molecule fluorescence microscopy by directly tracking rrnTAC assembly onto the functional TEC, transcription of rRNA and its processing by RNase III, all in a single experiment. Our data provide the first detailed mechanistic and quantitative understanding on how these events are functionally coupled by providing a molecular movie visualizing the entire process in real-time (Fig. 7): rrnTAC assembly is initiated by transient binding of NusA and NusG to RNAP - as it also would occur for mRNA transcription - and binding of NusE and NusB to the rRNA-specific boxBAC rRNA 5’-leader. SuhB binds last, stabilizing the entire complex. We show that only the fully assembled and stable complex is functional in accelerating transcription elongation and demonstrate an additional crucial function of the stably assembled rrnTAC: it dramatically increases the efficiency of co-transcriptional processing by RNase III, the step that marks the transition from co-transcriptional to post-transcriptional ribosome assembly.

**Figure 7.**
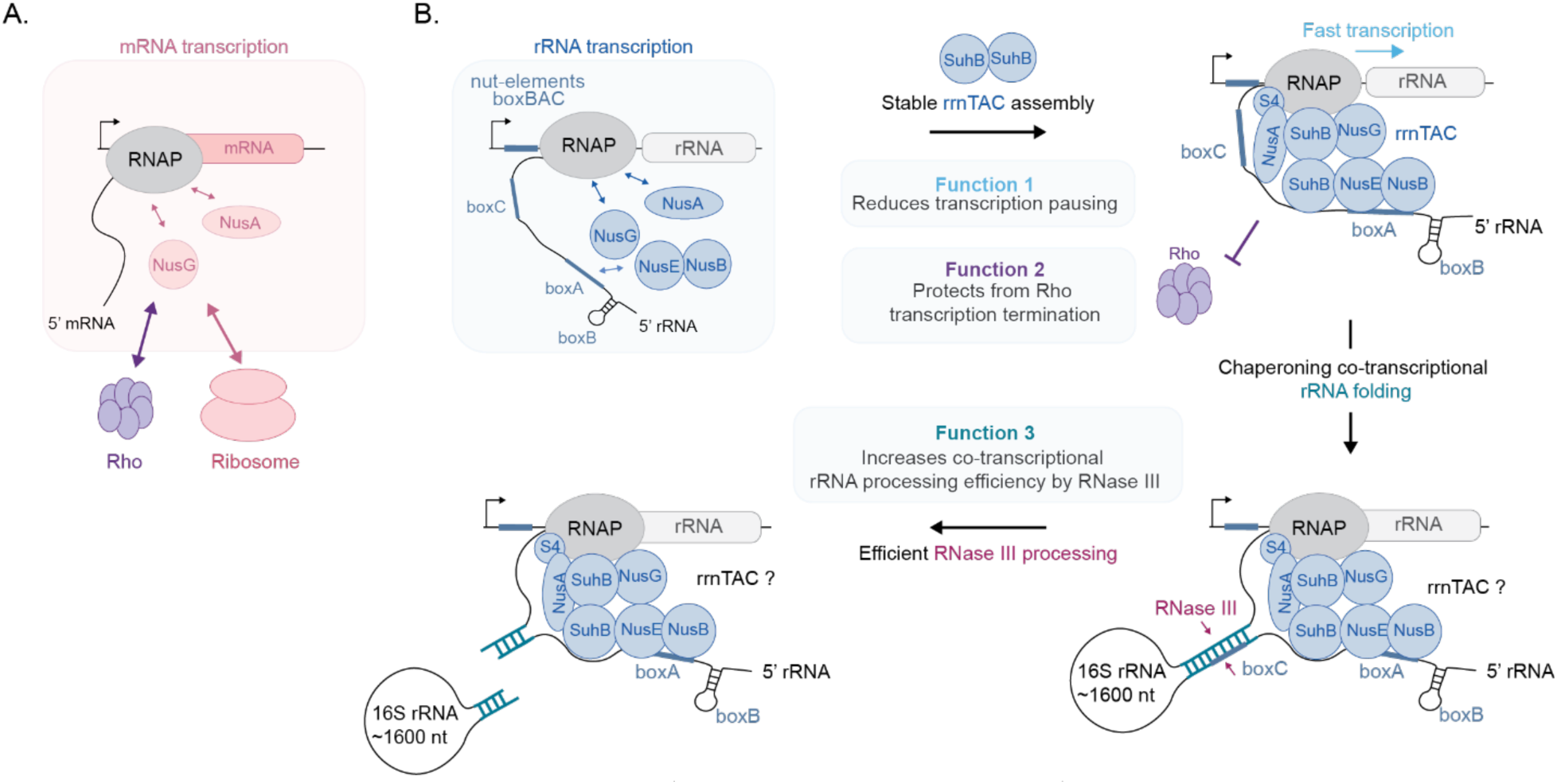
Model on how bacterial transcription factor dynamics determine function. (A) mRNA transcription: transient binding of NusA and NusG allows for regulation by Rho and the ribosome. (B) rRNA transcription: stable rrnTAC assembly is required for rRNA transcription and processing by RNase III.

### Transient versus stable association of transcription factors as a regulatory principle for rRNA versus mRNA transcription

CryoEM structures exist for NusA and NusG bound to the transcription elongation complex ^34^, representing the molecular composition of the complex required for mRNA transcription. Structures are also available of the TEC bound to all the rrnTAC proteins ^22^, representing the elongation complex for rRNA transcription. While these structural studies provide high-resolution snapshots, they cannot distinguish between functional versus non-functional interactions, they do not report on the underlying dynamics of the processes as they occur in real time, and crucially, they only provide limited information on how these processes may influence other processes simultaneously happening in a cellular context. Here, by quantifying the NusA and NusG residence times on the TEC (in presence of all the other rrnTAC factors), we find that NusA and NusG bind only for a few seconds (Fig. 2D) in absence of the rRNA-specific boxBAC sequence. However, NusA and NusG remain associated with the TEC for minutes (limited by photobleaching) if boxBAC is present (Fig. 3D). Thus, the binding dynamics of NusA and NusG serve fundamentally different biological purposes depending on the transcription context. During mRNA transcription, transient NusA/G binding enables rapid responsiveness to cellular signals: at high translational demand, NusG can simultaneously interact with the ribosome for functional coupling of transcription and translation, while in absence of ribosomes, NusG can interact and recruit Rho for premature transcription termination ^35–38^. Thus, dynamic transcription factor binding allows cells to immediately respond and couple mRNA synthesis to translation status. In contrast, rrnTAC assembly faces a different regulatory logic: once the complex has stabilized on RNAP through SuhB incorporation, regulation of transcription elongation becomes unnecessary because the rrnTAC serves to synthesize and process the entire rRNA transcript as efficiently as possible. Thus, stable rrnTAC assembly eliminates the need for dynamic regulation, contrasting sharply with the transient, regulatable state of NusA/G in mRNA contexts. This regulation mechanism reveals how the same conserved transcription factors (NusA, NusG) can adopt distinct functional states - transient and regulatable for mRNA, stable and committed for rRNA transcription (Fig. 7).

### rrnTAC assembly as double-specificity filter to prevent wasteful sequestration of factors

We show that rrnTAC assembly is initiated by weak and transient interactions with SuhB locking the entire assembly into a stable functional state, and the functional complex remaining stably assembled for minutes. Commitment to stable assembly is ensured by a double specificity filter: a boxBAC leader sequence engaging with NusE/B, thereby distinguishing rRNA from mRNA transcription (see above), and the binding of SuhB for stabilization (possibly regulated by rrnTAC protein levels: see below). This double filter is important for the following reasons: first, non-functional but stable assembly would sequester the rrnTAC proteins, depleting the pool of free transcription factors available for productive rRNA synthesis. Second, stable sequestration of processed boxBAC RNA by NusE/B would result in high and likely toxic accumulation of boxBAC RNA (>1000 copies generated per minute)Ehrenberg, et al. ^12^, Gotta, et al. ^13^. Third, early and premature stable engagement of proteins with the nascent boxBAC RNA could stabilize misfolded or non-native nascent RNA conformations incompetent of assembly into a functional complex within a biologically relevant time frame. Indeed, this assembly strategy, namely building a stable complex from multiple initial weak interactions, is reminiscent of r-protein assembly on nascent rRNA ^4,5,10^, and thus appears to be a general principle for coordinating rapid co-transcriptional assembly under the time constraints imposed by active transcription.

### Cellular rrnTAC protein levels may provide regulation by tuning rrnTAC assembly efficiency

We find a critical role of NusB and NusE in enabling efficient SuhB recruitment to the boxBAC-containing TEC and can provide a mechanistic explanation for previously puzzling *in vivo* observations: *nusB* defective mutants are cold-sensitive, but viable ^31,39^. This is consistent with our finding that loss of NusB dramatically slows SuhB recruitment (from ∼5 seconds to ∼100 seconds) but eventually still allows stable rrnTAC assembly, with that stable assembly being crucial for co-transcriptional RNase III processing.

In absence of successful SuhB recruitment, the rrnTAC remains in a transient, non-functional state. Thus, SuhB may also constitute a potential antibiotic target: pharmaceutical interference with SuhB recruitment would render rrnTAC complexes short-lived (few seconds lifetime) and non-functional.

### rrnTAC chaperones rRNA by both increasing transcription speed and by stably bridging 5’/3’-rRNA ends

Ribosome assembly requires co-transcriptional binding of r-proteins, with efficiency of assembly being challenged by possible rRNA misfolding which would mask binding sites of r-proteins. Local misfolding of RNA structures can also sequester RNA strands hampering the co-transcriptional formation of long-range RNA interactions ^10,40^. Ribosomal proteins themselves act as chaperones to fold rRNA ^4,5^ and DEAD-box helicases help to resolve kinetically trapped intermediates while not affecting stable protein-RNA complexes ^41^. Our data show another mechanism for chaperoning co-transcriptional rRNA folding: by physically and stably bridging the boxBAC leader with the RNAP, creating an RNA loop, the stably assembled rrnTAC maintains proximity between the 5’ and 3’ halves of the RNase III substrate helix throughout transcription (∼20 seconds for the 17S rRNA). Even brief dissociation of the 5’ half from the RNAP would allow the nascent 5’ end to engage in non-native structures, which form on the micro- to millisecond time scale as the nascent RNA emerges from the RNAP. In addition, the rrnTAC accelerates transcription speed approximately 2-fold (Fig. 1D, Fig. S7). Increased transcription speed reduces the time window during which the nascent rRNA can engage in non-native interactions ^4^ and this model is consistent with our observations of increased RNase III processing at higher transcription rates (Fig. 6H, and Fig, S7). Thus, rrnTAC chaperoning operates through two synergistic mechanisms: maintaining physical 5’-3’ proximity through rRNA looping and minimizing the time-window during which misfolding can occur, achieved by transcription acceleration.

### S4 is dispensable for the core rrnTAC functions

Among the six rrnTAC proteins, S4 occupies a unique position. Unlike all the other factors, S4 is not required for the core rrnTAC functions that we tested in our assays: presence of S4 did not increase transcription elongation rates (Fig. 1D, Fig. S1B), Rho-dependent transcription termination in presence of the other rrnTAC proteins (Fig. 1E, S1B), assembly speed (Fig. S5C) and stability (Fig. S5B) of the rrnTAC, nor co-transcriptional processing efficiency by RNase III (Fig. S9D).

These findings raise the question why S4 is associated with the rrnTAC. Although physically associated with the rrnTAC complex, its precise conformation remains unresolved in the cryoEM structure because of its flexibility ^22^. Due to its positioning close to the rRNA exit tunnel, S4 was suggested to act as an RNA-annealing factor by transiently interacting with the emerging RNA, possibly influencing its local RNA structure. While we did not observe an effect of S4 on RNase III processing efficiency, which reports on long-range helix formation, we cannot exclude the additional contribution of local chaperoning effects not detectable in our assays. Furthermore, r-protein S4 association with the rrnTAC could provide a mechanism to facilitate its co-transcriptional delivery to the 5’ domain of the 16S rRNA, where S4 acts as primary binding protein nucleating r-protein binding to the nascent rRNA. Binding of the primary r-protein S4 to the nascent rRNA is inefficient *in vitro* ^4,5^. By transiently traveling with the transcription machinery, S4 could immediately be delivered once its binding site on the rRNA has emerged from the RNAP, thereby seeding initiation of co-transcriptional ribosome assembly.

### RNA looping may provide a general mechanism for coordinating transcription-coupled processes

Beyond the specific context of bacterial co-transcriptional rRNA processing, our findings point to co-transcriptional RNA looping, in this specific case maintained by stable rrnTAC-mediated bridging of 5’ and 3’ rRNA ends, as a potentially universal strategy for eukaryotic and prokaryotic cells to coordinate multiple processes linked to transcription. For example, we recently demonstrated that ribosomes can reactivate paused RNAP through mRNA looping ^38^, thereby allowing long-range communication between the bacterial transcription and translation machineries. In eukaryotes, the U1 snRNP interacts with RNA polymerase II ^42^. If this interaction persists during transcription of long introns, 5’SS and 3’SS would be brought into proximity and provide an elegant mechanism for efficient co-transcriptional splicing of long introns, thus, coordinating transcription and splicing by RNA looping. The selection of mRNA 3’ ends is influenced by their transcription start site choice ^43,44^, a process which is mechanistically not yet understood but may also be coordinated by co-transcriptionally bridged 5’ and 3’ ends, mediated by Pol-II and RNA binding proteins. It remains to be investigated whether RNA looping is broadly functionally exploited in cells to coordinate processes linked to transcription.

### Limitations of the study

Although we believe that our complex *in vitro* reconstitutions in this study, and the simultaneous tracking of multiple cellular processes in real-time, provides the most complete system to date, several technical limitations remain to entirely reproduce the *in vivo* situation in a test tube. In presence of the completely assembled rrnTAC, the observed RNase III cleavage efficiencies are 56 % and 34 % for the 5’ domain and the complete native 17S rRNA, respectively, while *in vivo* the efficiencies are expected to be close to 100 %. Our minimal reconstitutions lack several cellular factors and do not occur in the same environment as in a living cell. Apart from missing r-proteins and cellular assembly factors, such as RNA modification enzymes and RNA chaperones ^2,3,41^, the high local protein concentrations or biomolecular condensates, in which bacterial ribosome assembly was recently suggested to occur ^45,46^ may be required for optimal rRNA folding and processing. The different environment could also explain our observation of a median rrnTAC assembly time of ∼5 seconds, while *in vivo* this may occur within 1-2 seconds. Nevertheless, our *in vitro* data provides mechanistic explanations for known *in vivo* phenotypes, such as the 20-fold reduced rrnTAC assembly speed without NusB (kinetics of SuhB recruitment: median 5 seconds with all factors, 110 seconds without NusB). Furthermore, our rates should be interpreted as intrinsic rates of the core machineries, rather than representing *in vivo* dynamics, which emerge from a complex convolution by the combined set of cellular factors under native conditions.

As generally true for all single-molecule fluorescence experiments in which the complexes of interest are surface-immobilized to enable minutes-long tracking of complex processes in real-time, the temporal resolution (∼ 100 ms per frame) limits the detection of interactions with dwell times shorter than ∼ 100 ms. Thus, transient encounters of the rrnTAC proteins with the transcription elongation machinery may be missed, even though they may still contribute productively to help rRNA folding and processing *in vivo* ^4,5^.

Our minimal reconstitution system demonstrates the intrinsic requirement for a stable rrnTAC for co-transcriptional RNase III processing of the rRNA (3% cleavage without SuhB versus >30% cleavage with stable rrnTAC assembly). However, it does not capture compensatory pathways that are present in cells. While mutations in the leader/trailer helix render ribosomes completely unfunctional ^47^, mutations in boxA reduce translation activity only 2-3-fold ^47^. Furthermore, RNase III processing of rRNA is not essential and can be rescued by other RNA processing enzymes at a highly decreased rate ^48^. The strong dependence of co-transcriptional processing on a fully assembled rrnTAC observed in our study demonstrates the intrinsic capacity of the rrnTAC to chaperone co-transcriptional rRNA processing, an effect only identifiable in a minimal active reconstitution. The reduced phenotypes of rrnTAC loss *in vivo* reflect cellular robustness, where backup mechanisms support crucial processes, rather than a minor role for the rrnTAC. Consequentially, pharmacologically targeting the rrnTAC may provide a robust way to inhibit ribosome assembly *in vivo*, if appropriately combined with suppressing the compensatory mechanisms.

## Acknowledgements

We thank Martyn Reynolds and Felix Jan Evers from Cairn Research/ Ultimeyes, Kavan Gor, Andrey Revyakin, Eva-Maria Geissen, the EMBL Chemical Synthesis Core Facility, EMBL Advanced Light Microscopy Facility, the EMBL IT team and Thomas Hoffmann for help and input in setting up the multi-color TIRF microscope and data evaluation pipeline. We also thank the EMBL Protein Expression and Purification Core Facility for purification of recombinant *E. coli* RNA polymerase. We thank Karine Lapouge for assistance in mass-photometry experiments. We would like to thank Benjamin Lau and Sernur Sena Yildiz for sharing NusE and Rho protein batches, respectively. We thank the entire Duss lab for helpful discussions. We would like to thank Benjamin Lau and Julia Mahamid for critically reading our manuscript. O.D. acknowledges support from the FEBS Excellence Award, the European Molecular Biology Laboratory and Deutsche Forschungsgemeinschaft (DFG project number 512397425).

## Author contributions

Conceptualization, A.C. and O.D.; methodology, A.C. and O.D.; Investigation, A.C. and N.Q.; writing—original draft, A.C.; writing—review & editing, A.C., N.Q. and O.D.; funding acquisition, O.D.; resources, O.D.; supervision, O.D.

## Supplementary information

Document S1. Figures S1–S10, Tables S1, S2 and S3, supplementary material and supplemental references

## Materials and methods

### Cloning

Wild-type sequences of NusA, NusB, NusG, NusE, SuhB, S4, RNase III and Rho were obtained from the ASKA plasmid collection ^49^ and cloned into a peSUMO vector backbone with Gibson assembly. For subsequent fluorescent labeling, single cysteine mutants of NusA (C251S_C454S_C489S_S53C and C251S_C454S_C489S_K243C), NusG_S60C, NusE_G103C and SuhB_C178S_S69C were constructed using site-directed mutagenesis ^50^. For RNase III, a ybbR-tag (DSLEFIASKLA) ^51^ was incorporated at the N-terminus. For RNAP, the ybbR-tag was inserted between E15 and E16 in the β′ subunit as described previously ^38^.

### Protein expression and purification

NusB, NusE, NusA, NusG, SuhB, S4, RNase III, Rho and respective mutans were overexpressed as N-terminal fusions with His_6_–SUMO. WT NusA, WT NusG and their mutants were purified as previously described ^38^. WT NusB, WT SuhB, SuhB mutants and WT Rho were purified using the same protocol with few modifications. In brief, cells were first lysed in IMAC buffer: 50 mM Tris-HCl pH 8.0, 300 mM NaCl (150 mM for NusB), 10-20 mM imidazole (no imidazole for WT NusB, WT SuhB and SuhB mutants), 10 mM 2-mercaptoethanol. After centrifugation, the lysate was loaded to a pre-equilibrated 5 ml HisTrap HP column (Cytiva) and eluted with increasing imidazole concentrations over 20 column volumes. Protein fractions were pooled together and dialyzed overnight against IMAC buffer (heparin buffer A for WT Rho: 20 mM Tris-HCl pH 8.0, 100 mM NaCl, 0.5 mM EDTA, 1 mM DTT) with 2 µM Ulp1 protease to cleave the fusion tag. A second HisTrap purification was done to remove the cleaved tag (except for WT Rho). WT Rho was loaded on a pre-equilibrated (with heparin buffer A) 5 ml HiTrap Heparin HP column (Cytiva) and eluted with increasing NaCl concentration (1 M NaCl in the final buffer). Protein fractions were then pooled, concentrated using 15 ml Amicon Ultracel 10 or 30 K concentrators. Next, the sample was purified further via size exclusion chromatography using either HiLoad S75 (WT NusG and NusG mutants) or HiLoad S200 (WT NusB, WT NusA and NusA mutants, WT SuhB and SuhB mutant, WT Rho) columns (Cytiva) equilibrated with storage buffer (20 mM Tris-HCl pH 7.6, 100 mM NaCl (50 mM for WT NusB, 200 mM for WT SuhB and SuhB mutants), 0.5 mM EDTA/DTT). The desired fractions were pooled and concentrated using 15 ml Amicon Ultracel 10 or 30 K concentrators. Aliquots were flash-frozen with liquid nitrogen and stored at −80 °C.

For NusE purification (WT and the mutant), the cells were lysed in buffer C (50 mM Tris-HCl, pH 7.5, 400 mM NaCl, 7 mM 2-mercaptoethanol) and centrifuged to pellet inclusion bodies. The pellet was washed twice with IB buffer (20 mM Tris-HCl pH 7.5, 100 mM NaCl, 0.5% (v/v) Triton X-100) by gently resuspending the pellet. Next, the pellet was solubilized in buffer containing 50 mM Tris-HCl pH 7.5, 6 M guanidine hydrochloride and centrifuged to remove the insoluble fraction and dialyzed overnight against buffer C (+2 M urea). The following day, the protein was centrifuged to remove the precipitated fraction and loaded on a pre-equilibrated 5 ml HisTrap HP column (Cytiva) with buffer C (+2 M urea) and eluted with increasing imidazole concentrations over 16 column volumes (to 0.5 mM imidazole in the final buffer). Fractions were analyzed on an SDS-PAGE gel and the desired fractions were pooled together and dialyzed in buffer C (+2 M urea) overnight with 2 µM Ulp1 protease. The following day, the protein was centrifuged to remove the precipitated fraction and loaded on a pre-equilibrated 5 ml HisTrap HP column (Cytiva) with buffer C (+2 M urea) for the second round of purification to remove the cleaved tag and the flow-through fractions were collected. Desired fractions were pooled and concentrated using a 15 ml Amicon Ultracel 3K concentrator. Aliquots were flash-frozen with liquid nitrogen and stored at −80 °C.

S4 purification was adapted from a previously described protocol ^52^. In brief, cells were lysed in buffer B (50 mM Tris/HCl pH 7.6, 400 mM KCl, 1 M NH_4_Cl, 7 mM 2-mercaptoethanol). The clarified lysate was loaded on the pre-equilibrated 5 ml HisTrap HP column (Cytiva) and eluted with increasing imidazole concentrations over 12 column volumes (to 0.5 M imidazole in the final buffer). Then, the desired fractions were pooled together and dialyzed overnight against buffer B with 2 µM Ulp1 protease. Then the second round of HisTrap purification was done. Desired fractions were pooled together and dialyzed overnight against SP buffer (20 mM Tris-HCl pH 7.0, 20 mM KCl, 6 M urea, 6 mM 2-mercaptoethanol) and loaded on a pre-equilibrated 5 ml HiTrap SP FF column (Cytiva) and eluted with increasing KCl (20 mM-350 mM KCl) concentration over 10 column volumes. Protein fractions were pooled together and dialyzed overnight against the storage buffer 50 mM Tris-HCl pH 7.6, 1 M KCl, 7 mM 2-mercaptoethanol. The sample was concentrated using a 15 ml Amicon Ultracel 10K concentrator and aliquots were flash-frozen with liquid nitrogen and stored at −80 °C.

The RNase III N-ybbR mutant was purified as previously described ^53^ with some modifications. In short, cells were lysed in buffer A (20 mM Tris-HCl pH 7.9, 500 mM NaCl, 5 mM imidazole, 1 mM 2-mercaptoethanol). The sample was purified using Ni-NTA agarose (Thermo Fischer) beads. Typically, 2 ml beads were used for purification of 5 grams of cell pellet. Clarified lysate was loaded onto the pre-equilibrated beads with buffer A. The beads were washed with 10 column volumes of buffer A, 6 column volumes of buffer A (+ 60 mM imidazole). The sample was eluted with elution buffer (20 mM Tris-HCl pH 7.9, 1 M NaCl, 400 mM imidazole, 1 mM 2-mercaptoethanol) over 4 column volumes and 0.5 ml fractions were collected. Fractions were analyzed on an SDS-PAGE gel and the desired fractions were pooled and dialyzed overnight against buffer A containing 2 µM Ulp1 protease to remove the cleaved tag. Another round of purification using Ni-NTA agarose (Thermo Fischer) was performed, and the flow-through fractions were pooled together and dialyzed overnight against the final buffer containing 30 mM Tris-HCl pH 8.0, 500 mM NaCl, 0.5 mM EDTA, 0.5 mM DTT. Sample aliquots were flash-frozen with liquid nitrogen and stored at −80 °C.

### Protein fluorescence-dye labeling

Single cysteine mutants of NusA, NusG, NusB, NusE, SuhB were labeled with sulfo-Cy5 (Lumiprobe) or sulfo-Cy5.5-maleimide (Cytiva) fluorophores. First, 0.5 mg or 1 mg of protein was reduced for 2 hours on ice in labeling buffer A (100 mM Na_2_HPO_4_/KH_2_PO_4_, 100 mM NaCl) with 10 mM DTT. Next, proteins were precipitated with 70 % (w/v) ammonium sulfate, centrifuged for 20 min at 14000 rpm after 20 min incubation and washed twice with ice-cold labeling buffer A with 70 % (w/v) ammonium sulfate. The protein pellets were dissolved in labeling buffer A (+6 M urea for NusB) and then incubated with 1 mg of DMSO dissolved fluorescent dye for 30 min at room temperature and shaking at 50 rpm. The labeling reaction was quenched with 0.5 % 2-mercaptoethanol. To remove the excess dye, the protein solution was loaded on a pre-equilibrated NAP-5 column (Cytiva) and eluted using labeling buffer A (+6 M urea for NusB). Collected fractions were analyzed with SDS-PAGE followed by fluorescent imaging. To perform the second round of purification, fractions containing labeled protein (except of NusB) were pooled together and loaded on a pre-equilibrated NAP-5 column, followed by analysis of the fractions on an SDS-PAGE gel and fluorescent imaging. For NusB labeling, the protein was refolded by a 1:10 dilution in heparin buffer A (20 mM Tris-HCl pH 7.6, 20 mM NaCl, 0.5 mM EDTA) and loaded onto a pre-equilibrated 1ml HiTrap Heparin HP column (Cytiva) with heparin buffer A and eluted with heparin buffer B (20 mM Tris-HCl pH 7.6, 1 M NaCl, 0.5 mM EDTA). Fractions were analyzed by an SDS-PAGE gel, followed by fluorescent imaging. Sample aliquots were flash-frozen with liquid nitrogen and stored at −80 °C.

To label the RNase III-ybbR mutant, 10 µM of protein was mixed with equal molar amounts of recombinantly purified SFP and 40 µM CoA-Cy5 dye in buffer containing 30 mM Tris-HCl pH 8.0, 0.5 M NaCl, 10 mM MgCl_2_, 2 mM DTT. Reactions were incubated for 3 hours at room temperature, then centrifuged for 5 min at 25000 g and loaded on a pre-equilibrated Zeba Spin desalting column (7K MWCO) (Thermo Fisher) with 30 mM Tris-HCl pH 8.0, 500 mM NaCl, 0.5 mM EDTA, 1 mM DTT. After two rounds of purification, the sample was analyzed by an SDS-PAGE gel, followed by fluorescent imaging. Sample aliquots were flash-frozen with liquid nitrogen and stored at −80 °C.

N-terminally tagged ybbR RNAP mutant was labeled with Cy5 as described previously ^38^. In short: 7 µM RNAP was mixed with 14 µM SFP, 28 µM CoA-Cy5 dye in a buffer containing 50 mM HEPES-KOH pH 7.5, 50 mM NaCl, 10 mM MgCl_2_, 2 mM DTT and 10% (v/v) glycerol ^54^. Labeling reactions were incubated at 25 °C for 2 h and analyzed on an SDS-PAGE gel, followed by fluorescent imaging. Sample aliquots were flash-frozen with liquid nitrogen and stored at −80 °C.

### Mass photometry

Mass photometry experiments were performed on a RefeynMP. The SuhB-Cy5 protein was diluted to 333 nM in degassed and filtered buffer containing 50 mM Tris pH 7.5, 3.5 mM MgCl_2_, 20 mM NaCl, 150 mM KCl, 0.04 mM EDTA, then 1 μl of the sample was added to 19 μl of the same buffer residing on the sample carrier slide and mixed by gentle pipetting. Particle mass was detected as the interference of scattered light with the surface-reflected light. Mass calibration was performed by using BSA.

### In vitro transcription and splinted ligation

RNA *in vitro* transcriptions were performed from PCR generated templates with sequences summarized in Table S1. Constructs were all containing a T7 promoter (TAATACGACTCACTATAG), a sequence for subsequent immobilization of the RNA to the glass surface for single-molecule imaging (AC-rich sequence: ACTACCACCACCCAACCAACACACC) and a sequence for labeling with a fluorescently labeled DNA oligonucleotide (F2 sequence: AACCACTCCAATTACATACACC), either inserted at the 5’end or at the 3’end of the RNA sequence of interest. For rnaAC015-rnaAC017 encoding the substrate helix for RNase III cleavage assays (see Table S1, Fig. 5), the p0030 sequence (TCACGAAAGCTGAGTAGTCACGAGTCTTCT) for labeling with the second fluorescent oligo was placed at the 5’end of the RNA, and the F2 sequence was placed in the loop region of the RNA. For RNAs used for scaffold assembly (rnaAC001, Table S1) or for splinted ligation (rnaAC011, Table S1), a minimal stem loop recognized by the VS ribozyme was placed at the 3’ end, allowing generating transcripts with homogeneous 3’ ends ^55^.

Transcription was performed using 30 nM DNA template in 40 mM Tris-HCl pH 8.0, 20 mM MgCl_2_, 1 mM spermidine, 0.01% Triton X-100, 5 mM of each NTP, 5 mM DTT and 1.7 µM in-house generated T7 RNAP for 4 hours at 37 °C. For reactions with VS-ribozyme cleavage, the reactions contained also the plasmid encoding the VS ribozyme. The VS ribosome plasmid was linearized by HindIII (NEB) restriction digestion overnight at 37 °C and was added to the transcription reaction at a final concentration of 2 nM with increased MgCl_2_ concentrations of 50 mM. Reactions were stopped by adding EDTA to a final concentration of 0.1 M and subjected to ethanol precipitation. After precipitation, the RNA was dissolved in nuclease free water and RNA loading dye (7 M urea in 2xTBE with 50 mM EDTA) was added in 1:1 ratio. RNA products were resolved on a 6 M urea PAGE gel (10 %; or 6 % for reactions with VS ribozyme cleavage). RNA bands were cut and purified using the crush and soak method ^56^, followed by ethanol precipitation and dilution in nuclease-free water. For RNAs generated with VS-ribozyme cleavage, the 2’-3’-cyclic phosphate was resolved by T4 polynucleotide kinase treatment in 1x PNK buffer for 4 hours at 37 °C, followed by enzyme inactivation (20 min at 65 °C), phenol-chloroform extraction and ethanol precipitation. The final RNA was dissolved in nuclease free water.

Splinted ligation of the two RNAs (rnaAC011 and rnaAC012-Cy3, Table S1) was performed by first annealing both RNAs to the splint DNA oligonucleotide (prAC197, Table S3) at 14 µM concentration in 1 x e55 buffer (10 mM Tris-HCl pH 7.5, 20 mM KCl) for 5 min at 95 °C, followed by a slow-cool down until room temperature. Next, the ligation reaction was performed at 8.4 µM concentration in splint ligation buffer (40 mM Tris-HCl pH 7.8, 10 mM MgCl_2_, 0.5 mM ATP, 10 mM DTT) with 4 µM T4 DNA ligase for 2 hours at 37 °C ^55^. The RNA loading dye was added to stop the reaction in 1:1 ratio and the sample was heated for 2-3 min at 95 °C and resolved on an 10 % PAGE (6 M urea) gel. The ligated RNA was cut out from the gel and extracted using the crush and soak method ^56^ with subsequent ethanol precipitation and final dilution in nuclease free water.

### DNA connectors and labeling

Different linear DNA sequences (connector DNA) (Table S2) were generated by PCR either from a purchased gene block (IDT) or from the pKK3535 plasmid encoding the native *rrnB* operon. Additionally, the BsaI restriction site was introduced at the 5’ end of the generated fragment. The 3’end of the DNA fragment encoded T7Te sequence (TAATCACACTGGCTCACCTTCGGGTGGGCCTTTCTGCGTTTAT), followed by a single stranded overhang (GGGATGGTAATTTGGTGAGTATGATTAAGGatctcCTCGAGgactagctg) introduced by autosticky PCR ^57^. PCR products were purified using a PCR purification kit (QIAGEN), and cleaved overnight with BsaI (NEB) at 37 °C, followed by enzyme inactivation for 15 min at 80 °C. Next, connector DNA was purified again using a PCR purification kit (QIAGEN), re-buffered to the storage buffer (10 mM Tris-HCl pH 7.5, 20 mM KCl) using an Amicon concentrator of 30K MWCO (Sigma). Next, the 3’ end single-stranded overhang was hybridized to the two DNA oligos (p0088–2xCy3.5, prAC143_biotin) (Table S3), which was added in an 1.2-fold excess in scaffold buffer (20 mM Tris-HCl pH 7.5, 40 mM KCl, 1 mM MgCl_2_) and incubated 5 min at 65 °C, followed by slow-cool down at room temperature. Hybridization of the 3’end overhang to the prAC143_biotin oligo generated a double-stranded XhoI restriction site (Table S2).

### Scaffold assembly and purification

*In vitro* transcribed RNA, template DNA (tDNA) and non-template DNA (ntDNA) were annealed at 1.2 µM (0.4 µM for scaffold assembly of γ-[^32^P]-RNA) in scaffold buffer (20 mM Tris-HCl pH 7.5, 40 mM KCl, 1 mM MgCl_2_) and incubated 5 min at 85 °C followed by a slow cool-down. Next, 500 nM of RNA:DNA scaffold (400 nM for scaffold with γ-[^32^P]-RNA) was mixed with 4 (3.5 for scaffold with γ-[^32^P]-RNA) equivalents of the RNAP core enzyme in transcription buffer (50 mM Tris-HCl pH 8, 20 mM NaCl, 14 mM MgCl_2_, 0.04 mM EDTA, 40 µg/ml non-acylated BSA, 0.01% (v/v) Triton X-100 and 2 mM DTT) and incubated for 20 min at 37 °C forming a partial scaffold (Table S2, Fig. S3A). The 5’end of the tDNA contained a phosphate group and together with the 3’end of ntDNA was designed to mimic the sticky ends generated by the BsaI restriction enzyme. To form a full-length scaffold, the partial scaffold was ligated to the fluorescently labeled connector DNA at a final concentration of 145 nM in 1xT4 ligase buffer (+20 mM KCl) with 1.25 µM in-house generated T4 DNA ligase (30 min, RT). Full-length scaffolds were purified using magnetic Streptavidin beads (NEB) using biotin at the 3’end of the connector. Beads (15 µl) were washed three times with 100 µl 1xPM2 buffer (50 mM TrisOAc (pH 7.5), 5 mM NH_4_OAc, 0.5 mM Ca(OAc)_2_, 5 mM Mg(OAc)_2_, 0.5 mM EDTA, 100 mM KCl, 1 mM spermidine, 5 mM putrescine). The ligation mixture was incubated with the beads in 1xPM2 buffer and incubated at room temperature for 30 min with frequent pipetting. Next, the beads were washed three times with 100 µl of 1xPM2 buffer (+250 mM KCl). For elution, beads were resuspended in transcription buffer (50 mM Tris-HCl pH 8, 20 mM NaCl, 14 mM MgCl_2_, 0.04 mM EDTA, 40 µg/ml non-acylated BSA, 0.01% (v/v) Triton X-100) followed by addition of XhoI (NEB) restriction enzyme and incubated for 20 min at 37 °C with frequent pipetting. The supernatant was collected and stored on ice.

### Bulk in vitro transcription and RNase III processing assays

Promoter-based single-round *in vitro* transcription assays were performed using a DNA template encoding the first 900 nucleotides of the pre-rRNA starting 30 bp upstream of P1 promoter and ending downstream of the 5’-domain followed by a T7Te sequence at the 3’end of the template (Table S2). The DNA was amplified with PCR. Stalled transcription elongation complexes (TECs) were formed in transcription buffer (50 mM Tris-HCl pH 8, 20 mM NaCl, 14 mM MgCl_2_, 0.04 mM EDTA, 40 µg/ml non-acylated BSA, 0.01 % (v/v) Triton X-100 and 2 mM DTT) incubating 25 nM DNA template (20 min at 37 °C) with 100 nM *E. coli* RNAP, 100 µM ACU trinucleotide, 5 µM GTP, 5 µM CTP, and 5 µM ATP (100 nM [α-^32^P]-ATP, Hartmann Analytic) thereby stalling RNAP at U34 and U43. To prevent re-initiation of transcription, rifampicin was added to the reaction at 10 µg/ml final concentration and incubated for 20 min at 37 °C. Next, 400 nM proteins (NusA only, NusG only, rrnTAC, rrnTAC without SuhB, rrnTAC without S4) were added to the TEC and incubated for 2 min at 37 °C. Transcription was re-started by addition of NTP chase mix (5 µM GTP, 5 µM CTP, 5 µM ATP, and 10 µM UTP) at 37 °C. Time points were taken before addition of the chase mix (“0”), and at 1 min, 2 min, 5 min, 10 min, and 30 min. The reaction was stopped by incubating 4 µl of sample with 4 µl stop buffer (7 M urea, 2xTBE, 50 mM EDTA, 0.025% (w/v) bromphenol blue and xylene blue) at 95 °C for 2 min. The reactions were analyzed on a 6 % denaturing PAGE gel (7 M urea in 1x TBE), running in 1x TBE at 50 W for 1 h 30 min, followed by gel drying for 1 h at 80 °C, overnight exposure on a phosphor screen (Cytiva, BAS IP MS 2040 E), and imaging using a Typhoon FLA 9500.

To perform bulk RNase III processing assays, 500 nM of substrate RNAs (rnaAC015, rnaAC016, rnaAC017; Table S1) were dephosphorylated using rSAP phosphatase (NEB) in T4 PNK buffer for 1 hour at 37 °C, followed by inactivation for 15 min at 65 °C. To radioactively label the 5’-end of the RNA with [γ-^32^P]-ATP, 200 nM of dephosphorylated RNA was mixed with equal amount of [γ-^32^P]-ATP (Hartmann Analytic) in T4 PNK buffer with T4 PNK (NEB) and incubated for 1 hour at 37 °C, followed by inactivation for 15 min at 65 °C. The labeled RNA was then purified with a G25 column (Cytiva) to remove not-incorporated [γ-^32^P]-ATP. Next, RNA was heated up to 85 °C for 3 min, cooled until room-temperature, and diluted to 20 nM with imaging buffer (50 mM Tris-HCl pH 7.5, 3.5 mM MgCl_2_, 20 mM NaCl, 150 mM KCl, 0.04 mM EDTA, 40 ug/ml BSA, 0.01 % Triton X-100, 2 mM spermidine, 1 mM putrescine). Reactions were initiated by addition of 20 nM unlabeled RNase III-ybbR mutant to RNA. Time points were taken before the addition of RNase III (“0”), and at 30 sec, 1 min, 2 min, 5 min, and 10 min. The reactions were stopped by incubating 15 µl of sample with 15 µl stop buffer (7 M urea, 2xTBE, 50 mM EDTA, 0.025 % (w/v) bromphenol blue and xylene blue) at 95 °C for 2 min. The reactions were analyzed on a 10 % denaturing PAGE (7 M urea in 1x TBE), running in 1x TBE at 50 W for 2 hours, followed by gel drying for 1 h at 80 °C, overnight exposure on a phosphor screen (Cytiva, BAS IP MS 2040 E), and imaging using a Typhoon FLA 9500.

To perform bulk co-transcriptional RNase III processing assays, 4.4 µM of rnaAC001 was dephosphorylated using rSAP phosphatase (NEB) in T4 PNK buffer for 1 hour at 37 °C, followed by inactivation for 15 min at 65 °C. To radioactively label the 5’-end of the RNA with [γ-^32^P]-ATP, 1.2 µM of dephosphorylated RNA was mixed with equal molar amounts of [γ-^32^P]-ATP (Hartmann Analytic) in T4 PNK buffer with T4 PNK (NEB) and incubated for 1 hour at 37 °C, followed by inactivation for 15 min at 65 °C. Labeled RNA was then purified with a G25 column (Cytiva) to remove not-incorporated of ^32^P γ-ATP. The scaffold was assembled as described in the previous section with partial scaffold 1 and connector (5’domain_spacer_T7Te) (Table S1-2). After scaffold purification, 10 nM of scaffold was incubated with 1 µM rrnTAC proteins (except S4, +/- SuhB) in imaging buffer (50 mM Tris-HCl pH 7.5, 3.5 mM MgCl_2_, 20 mM NaCl, 150 mM KCl, 0.04 mM EDTA, 40 ug/ml BSA, 0.01 % Triton X-100, 2 mM spermidine, 1 mM putrescine) for 10 min at 37 °C. The sample was chased with 0.2 mM NTPs +/- 50 nM RNase III and 6.5 mM MgCl_2_. Time points were taken before addition of chase mix (“0”), and at 10 sec, 20 sec, 40 sec, 1 min, 2 min, and 5 min. The reactions were stopped by incubating 10 µl of sample with 10 µl stop buffer (7 M urea, 2xTBE, 50 mM EDTA, 0.025 % (w/v) bromphenol blue and xylene blue) at 95 °C for 2 min. The reactions were analyzed on a 6 % denaturing PAGE gel (7 M urea in 1x TBE), running in 1x TBE at 50 W for 1 h 30 min, followed by gel drying for 1 h at 80 °C, overnight exposure on a phosphor screen (Cytiva, BAS IP MS 2040 E), and imaging using a Typhoon FLA 9500.

### Single-molecule instrumentation, acquisition and analysis

Single-molecule imaging experiments were performed on a custom-built objective based (CFI SR HP Apochromat TIRF 100×C Oil) TIRF microscope (built in collaboration with Cairn Research: https://cairn-research.co.uk/ and Ultimeyes: https://www.ultimeyes.eu/) equipped with an iLAS system (Cairn Research), three Prime95B sCMOS cameras (Teledyne Photometrics) and a diode-based (OBIS) 532 nm laser ^4,10,38^. The MetaMorph software package (Molecular Devices) was used for data acquisition. Single-molecule traces were extracted and processed from acquired movies using the SPARTAN software package (v.3.7.0) ^58^. Traces were further exported from SPARTAN and analyzed in MATLAB (version R2019a) ^4,10,38^. Further analysis in MATLAB included picking traces for evaluation (criteria specified for each experiment in the following) and analysis of protein-bound dwell times by assigning the FRET events using thresholding ^27,38^. Bound lifetimes were determined by fitting dwells with a single-exponential equation if not stated otherwise: y=1-exp(-k*t). In the following we describe the details for all recorded experiments.

Experiments to study binding of rrnTAC proteins to the stalled TEC lacking the boxBAC RNA elements (Fig. 2A-D): the stalled TEC was prepared as described previously ^4^ and the DNA template (T13, Table S2) was labeled with p088-2xCy3.5 (Table S3) (5 min at 65 °C with slow cooling to room temperature). The RNA in the stalled complex was labeled by hybridization with a Cy3-labeled DNA oligo (p066-Cy3, Table S3) 13 nt upstream from the RNAP exit channel and annealed to a biotinylated DNA duplex (p141p109-biotin, Table S3) allowing subsequent binding to the glass surface. The reaction was performed using 25 nM stalled TEC, 20 nM of biotinylated DNA duplex and 100 nM of p066-Cy3 oligo in transcription buffer by incubating for 20 min at 37 °C. The sample was diluted in the imaging buffer (50 mM Tris-HCl pH 7.5, 3.5 mM MgCl_2_, 20 mM NaCl, 150 mM KCl, 0.04 mM EDTA, 40 ug/ml BSA, 0.01 % Triton X-100, 2 mM spermidine, 1 mM putrescine) to 25 pM and immobilized on a biotin-polyethylene glycol functionalized glass slide ^59^. Prior to the immobilization, the glass surface was coated with NeutrAvidin for 5 min, followed by extensive washing with imaging buffer. The sample was incubated for 10 min and then washed with imaging buffer with addition of an oxygen scavenger system (OSC), containing 2.5 mM protocatechuic acid and 190 nM protocatechuate dioxygenase and a cocktail of triplet state quenchers (1 mM 4-citrobenzyl alcohol, 1 mM cyclooctatetraene and 1 mM Trolox) to minimize fluorescence instabilities. The reaction was initiated by injecting 50 nM Cy5-labeled rrnTAC proteins (except 10 nM for NusA-Cy5) in either absence or presence of 400 nM of all the other unlabeled rrnTAC proteins (100 nM for S4) at 37 °C. The NusA(C251S_C454S_C489S_K243C)-Cy5 mutant was used in single-molecule experiments if not stated otherwise. A 3-color set-up was used to detect Cy3, Cy3.5 and Cy5 signals simultaneously. The sample was excited with the 532 nm laser at 0.86 kW/cm^2^ (100%) output intensity and 3000 frames were acquired at 100 ms frame time. Molecules with both Cy3 and Cy3.5 signals (Cy3 signal present during the entire movie) with intensities corresponding to a single-molecule were selected for further evaluation in MATLAB.

Experiments to study the binding of the rrnTAC proteins to RNA encoding the boxBAC element (Fig. 2F-H): the 3’ end of the rnaAC002 RNA (Table S1) was hybridized to a biotinylated DNA duplex (p0155p0154-biotin, Table S3) for immobilization and to the p066_Cy3 DNA oligo (Table S3) for fluorescence labeling (5 min at 65 °C with slow cool down). The following steps for immobilization and washing were the same as described in the previous paragraph. The reaction was initiated by injecting 50 nM Cy5-labeled rrnTAC protein (10 nM for NusA-Cy5) in either absence (imaging was done at 37 °C) or presence of 400 nM of all the other unlabeled rrnTAC proteins (100 nM for S4) at 30 °C. A 2-color set-up was used to detect Cy3 and Cy5 signals simultaneously. The sample was excited with the 532 nm laser at 0.73 kW/cm^2^ output intensity and 3000 frames were acquired at 100 ms frame time. Molecules with a Cy3 signal present during the entire movie corresponding to the intensity of a single-molecule were selected for further evaluation in MATLAB.

Experiments to study binding of individual rrnTAC proteins (Fig. 3) or the co-binding of two labeled proteins at the same time to a stalled TEC containing the boxBAC element (Fig. 4F-I): the assembled and purified RNA:DNA scaffold consisting of partial_scaffold_1, rnaAC001 and RNAP core enzyme was first ligated to the prAC217 connector DNA (labeled with p088_2xCy3.5 oligo; Table S3), which is encoding the 5’domain_spacer_T7Te (Table S1-2). Then the entire assembly was hybridized to a DNA duplex (p141p109-biotin, Table S3) and to a Cy3-DNA oligo (p066-Cy3, Table S3) in transcription buffer for 20 min at 37 °C. Next, the sample was immobilized as described above. A single-molecule channel with two tube inlets was used for this type of experiment, allowing 2-step injection during the movie acquisition. The rrnTAC assembly experiment was initiated by injecting a first chase mix containing 50 nM of individual Cy5-labeled rrnTAC proteins (10 nM for NusA) either in absence or in presence of all the other rrnTAC factors unlabeled (or with a subset of factors) at 400 nM concentration (100 nM for S4) at 37 °C (data for NusB without rrnTAC, NusG without rrnTAC, SuhB without rrnTAC was measured at 30 °C). To score molecules for transcription activity, a second chase containing 200 µM NTPs was injected 6 minutes after the start of the experiment. The second injection allowed to select only transcriptionally active molecules for evaluation. A 3-color set-up was used to detect the Cy3, Cy3.5 and Cy5 signals simultaneously. The sample was excited with the 532 nm laser at 0.73 kW/cm^2^ output intensity, followed by 50 frames without laser excitation, and a final excitation with 532 nm laser at 0.73 kW/cm^2^ output intensity until the end of the movie. Injections were done during the time windows without laser excitation. A total of 6000 frames were acquired at 100 ms frame time. For the experiments with co-binding of two labeled proteins (Fig. 4F-I), we used 10 nM NusA-Cy5 (or 50 nM NusB-Cy5) and 50 nM SuhB-Cy5.5 with the other rrnTAC proteins unlabeled at 400 nM concentration (100 nM for S4). The molecules were scored for transcription activity with 200 µM NTPs 6 min after the start of the experiment. A 4-color set-up was used to detect the Cy3, Cy3.5, Cy5 and Cy5.5 signals simultaneously and the same pulse train as for 3-color set-up was used. For evaluation, we selected molecules with all the following criteria: 1) Cy3 signals corresponding to a single molecule intensity, 2) Cy3 signal present at least until the second injection, 3) Cy3.5 signal corresponding to active transcription after the second injection (identified by single-step drop of the Cy3.5 intensity as described previously ^27^.

Experiments to study RNase III processing of the pre-transcribed RNAs (Fig. 5): substrate RNAs (rnaAC015, rnaAC016, rnaAC017, Table S1) were hybridized with a Cy3-DNA oligo at the 5’ end (p066-Cy3, Table S3), a Cy3.5-DNA oligo (p0150_Cy3.5, Table S3) in the loop region of the RNA and a biotinlylated DNA duplex (p0155p0154-biotin, Table S3) in 10 mM Tris-HCl pH 7.5, 20 mM KCl for 5 min at 85 °C with slow cool down. The sample was diluted in the imaging buffer to 50 pM and was immobilized on the glass surface. After washing with OSC containing imaging buffer, imaging was started and 20 nM of RNase III - Cy5 (if not stated differently) in 3.5 mM MgCl_2_ (if not stated differently) was injected to initiate the reaction. A 3-color set-up was used to detect the Cy3, Cy3.5 and Cy5 signals simultaneously. The sample was excited with the 532 nm laser at 0.73 kW/cm^2^ output intensity, and 6000 frames were acquired at 100 ms frame time. Molecules containing both Cy3 and Cy3.5 signals corresponding to the intensity of a single-molecule which were present either until the RNase III - Cy5 cleavage or up to 3000 frames were selected for further evaluation in MATLAB.

Experiments to measure transcription speed (Fig. S7): the scaffolds were assembled using Cy5-labeled RNAP, rnaAC001, partial scaffold_1 and ligated to connector encoding the 16S rRNA directly followed by a T7Te terminator (prAC234) (Table S1-2, Fig. S3A). The connector DNA was labeled with the p088_2xCy3.5 oligo (5 min at 65 °C), and the RNA in the scaffold was hybridized to the p141p109-biotin oligo (20 min at 37 °C). To assemble the rrnTAC, 1 µM of each unlabeled rrnTAC protein (except S4) +/-SuhB in OSC imaging buffer was injected, and the sample was incubated for 10 min at 37 °C. Next, the chase mix containing different NTPs concentrations and 1 µM rrnTAC proteins (only for condition “without SuhB”) were injected in OSC imaging buffer 75 frames after the start of imaging. A 2-color set-up to detect the Cy3.5 and Cy5 signals simultaneously was used. The experiments were acquired with the following laser pulse sequence: 75 frames excited with the 532 nm laser at 0.73 kW/cm^2^ output intensity, followed by 15 frames without excitation, and excitation with the 532 nm laser at 0.73 kW/cm^2^ output intensity until 3000 frames at 200 ms frame time. Molecules showing Cy3.5 and Cy5 FRET prior to a single Cy3.5 intensity drop indicating the end of transcription ^38^ were selected for evaluation in MATLAB. The transcription time for each single molecule was defined from NTP injection till appearance of the RNAP-Cy5 / DNA-Cy3.5 FRET.

Experiments to track rrnTAC assembly and co-transcriptional RNase III processing (Fig. 6): the scaffold was assembled with internally labeled RNA (rnaAC013-Cy3, tDNA_prAC199, ntDNA_prAC198, Table S1-2) and ligated to either prAC217_5’domain_spacer_T7Te, or to prAC185_16SrRNA_spacer_T7Te (Table S2). After purification, the sample was hybridized to the DNA duplex (p141p109-biotin) in transcription buffer for 20 min at 37 °C. The final sample was diluted 100-fold and immobilized to the glass-surface via the 5’ end of the RNA (Fig. 6C). To assemble the rrnTAC, we introduced as the first chase mix 10 nM NusA-Cy5.5 (NusA C251S_C454S_C489S_S53C mutant used) with 1 µM of the other unlabeled rrnTAC proteins +/- SuhB (except S4) in OSC imaging buffer and incubated the sample for 10 min at 37 °C. To select molecules with stable binding of NusA-Cy5.5 (detected with FRET from the internal Cy3 dye on the RNA), we imaged the sample for 2 min. Then, to initiate rRNA transcription and processing by RNase III, we injected the second chase mix containing 0.2 mM NTPs (or 1 mM NTPs), 50 nM RNase III-Cy5, 10 mM MgCl_2_ in OSC imaging buffer. In the same chase mix we kept also the components of the first chase mix. To investigate the effect of S4 protein on co-transcriptional RNase III processing (Fig. S9D), the second chase mix contained 3 nM S4, 1mM NTPs, 50 nM RNase III-Cy5, 10 mM MgCl_2_ in OSC imaging buffer. A 4-color set-up was used to detect the Cy3, Cy3.5, Cy5 and Cy5.5 signals simultaneously. The experiment was acquired with the following laser pulse sequence: excitation with the 532 nm laser at 0.73 kW/cm^2^ output intensity for 600 frames, followed by 25 frames without excitation, and excitation with the same laser intensity until 3000 frames at 200 ms frame time. For evaluation in MATLAB, we selected molecules based on all the following criteria: 1) Cy3 intensity corresponding to the intensity of a single-molecule, 2) Cy3 signal present after injection, 3) Cy3.5 signal corresponding to active transcription after injection (selected by single-step drop of Cy3.5 intensity ^27^).

### Statistical analysis

Errors in dwell time fits represent 95 % confidence intervals obtained from fits to single-exponential functions as indicated. Analyzed molecules and number of dwells used in the dwell time analyses are described in the manuscript text, figure legends and supplementary information. For boxplots within the entire manuscript, median (central mark), 25th (bottom edge) and 75th (top edge) percentiles are indicated. Whiskers correspond to 1.5x interquartile range. **Data availability.** Structures were downloaded from the Protein Data Bank (https://www.rcsb.org/) using the accession codes shown in Fig. 1C and Fig. S2A. The published article includes all relevant data generated or analyzed during this study. The raw data files are too large (several TBs of data) but are available upon request to O.D. **Code availability.** The software used for single-molecule data processing and analysis is published ^58,60^ and freely available online. Additional scripts for downstream processing were made available publicly previously under an open-source license ^61^.

