## Supplementary Material for "Functional coupling between ribosomal RNA transcription and processing guided by stable transcription factor binding"

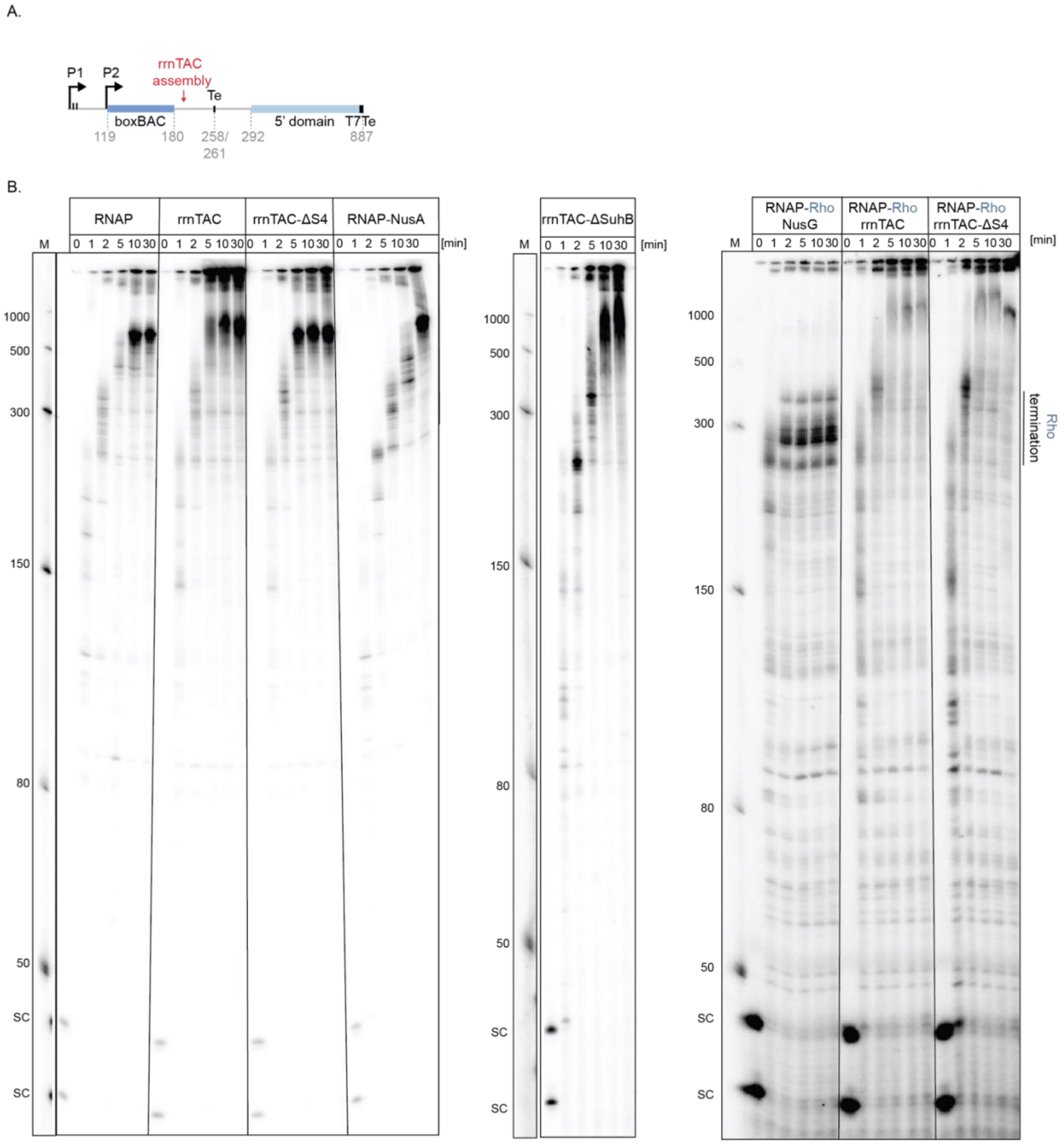


**Fig. S1. Reconstituted rrnTAC suppresses transcription pausing and Rho transcription termination.**

(A) Overview of the DNA template used for transcription assays with [α-$^{32}$P]-ATP. P1 and P2 indicate *rrnB* promoters. Two black lines after P1 show RNAP stalling positions (34 nt, 43 nt). BoxBAC elements are shown in blue, rrnTAC assembly point is marked with red and indicates the point when boxBAC elements are transcribed and out of the RNAP exit channel. Putative Rho-dependent terminator "Te" (described in [1]) is shown, followed by the 5' domain sequence and the T7Te terminator.

(B) Left gel: single-round transcription assays of RNAP without factors; 400 nM rrnTAC proteins; 400 nM rrnTAC proteins without S4; 400 nM NusA are shown. The stalled transcription elongation complex was formed with 25 nM DNA template, 100 nM *E. coli* RNAP, 100 μM ACU trinucleotide, 5 μM GTP, 5 μM CTP, and 5 μM ATP (100 nM [α-$^{32}$P]-ATP), thereby stalling (denoted as SC) RNAP at U34 and U43. Transcription was re-started by addition of NTPs chase mix (5 μM GTP, 5 μM CTP, 5 μM ATP, and 10 μM UTP) at 37 ºC. Time points were taken before addition of chase mix (“0”), and at 1 min, 2 min, 5 min, 10 min, and 30 min. Middle gel: Single-round transcription assays of RNAP with 400 nM rrnTAC proteins without SuhB. Reactions were performed in the same way as described for the left gel. Right gel: single-round transcription assay of RNAP in the presence of 100 nM NusG and 300 nM Rho (50 nM hexamer); 400 nM rrnTAC proteins and 300 nM Rho; 400 nM rrnTAC proteins omitting S4 and 300 nM Rho. Stalled transcription elongation complex was formed with 25 nM DNA template, 100 nM *E. coli* RNAP, 100 μM ACU trinucleotide, 5 μM GTP, 5 μM CTP, and 100 nM [α-$^{32}$P]-ATP stalling RNAP at U34 and U43. Transcription was re-started by addition of NTP chase mix (5 μM GTP, 5 μM CTP, 10 μM ATP, and 10 μM UTP) and 1 mM ATP for Rho activity at 37 ºC. Time points were taken before addition of chase mix (“0”), and at 1 min, 2 min, 5 min, 10 min, and 30 min. Parts of some gels are also shown in Fig. 1. Uncropped gels are shown in supplementary material file.

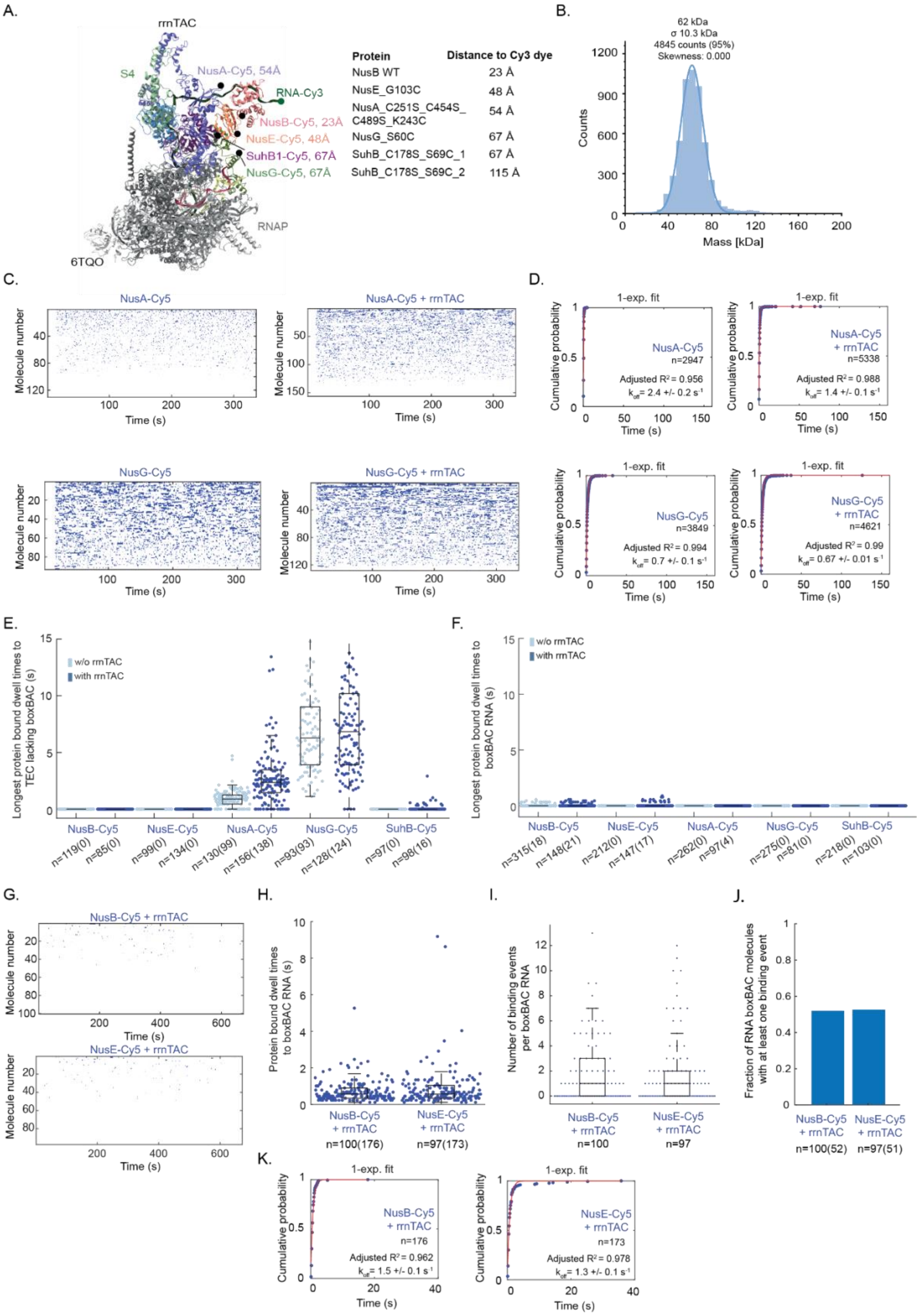
A.
rrnTAC
S4
NusA-Cy5, 54Å
RNA-Cy3
NusB-Cy5, 23Å
NusE-Cy5, 48Å
SuhB1-Cy5, 67Å
NusG-Cy5, 67Å
RNAP
6TQO
Protein | Distance to Cy3 dye
NusB WT | 23 Å
NusE_G103C | 48 Å
NusA_C251S_C454S_C489S_K243C | 54 Å
NusG_S60C | 67 Å
SuhB_C178S_S69C_1 | 67 Å
SuhB_C178S_S69C_2 | 115 Å
B.
62 kDa
σ 10.3 kDa
4845 counts (95%)
Skewness: 0.000
Counts
Mass [kDa]
C.
NusA-Cy5
NusA-Cy5 + rrnTAC
NusG-Cy5
NusG-Cy5 + rrnTAC
Molecule number
Time (s)
D.
1-exp. fit
Cumulative probability
NusA-Cy5 n=2947 Adjusted R² = 0.956 k_off = 2.4 +/- 0.2 s⁻¹
NusA-Cy5 + rrnTAC n=5338 Adjusted R² = 0.988 k_off = 1.4 +/- 0.1 s⁻¹
NusG-Cy5 n=3849 Adjusted R² = 0.994 k_off = 0.7 +/- 0.1 s⁻¹
NusG-Cy5 + rrnTAC n=4621 Adjusted R² = 0.99 k_off = 0.67 +/- 0.01 s⁻¹
Time (s)
E.
Longest protein bound dwell times to TEC lacking boxBAC (s)
w/o rrnTAC
with rrnTAC
NusB-Cy5 NusE-Cy5 NusA-Cy5 NusG-Cy5 SuhB-Cy5
n=119(0) n=85(0) n=99(0) n=134(0) n=130(99) n=156(138) n=93(93) n=128(124) n=97(0) n=98(16)
F.
Longest protein bound dwell times to boxBAC RNA (s)
w/o rrnTAC
with rrnTAC
NusB-Cy5 NusE-Cy5 NusA-Cy5 NusG-Cy5 SuhB-Cy5
n=315(18) n=148(21) n=212(0) n=147(17) n=262(0) n=97(4) n=275(0) n=81(0) n=218(0) n=103(0)
G.
NusB-Cy5 + rrnTAC
NusE-Cy5 + rrnTAC
Molecule number
Time (s)
H.
Protein bound dwell times to boxBAC RNA (s)
NusB-Cy5 + rrnTAC n=100(176)
NusE-Cy5 + rrnTAC n=97(173)
I.
Number of binding events per boxBAC RNA
NusB-Cy5 + rrnTAC n=100
NusE-Cy5 + rrnTAC n=97
J.
Fraction of RNA boxBAC molecules with at least one binding event
NusB-Cy5 + rrnTAC n=100(52)
NusE-Cy5 + rrnTAC n=97(51)
K.
1-exp. fit
Cumulative probability
NusB-Cy5 + rrnTAC n=176 Adjusted R² = 0.962 k_off = 1.5 +/- 0.1 s⁻¹
NusE-Cy5 + rrnTAC n=173 Adjusted R² = 0.978 k_off = 1.3 +/- 0.1 s⁻¹
Time (s)

**Fig. S2. Binding dynamics of rrnTAC proteins to the isolated TEC at 37 ºC and boxBAC RNA at 21 ºC.**

(A) Structure of rrnTAC/TEC complex (PDB: 6TQO) showing the labeling positions for each rrnTAC protein (black circle; acceptor dye) and the distance to the 5' end of RNA (green circle; donor dye).
(B) Mass-photometry showing complete SuhB homodimer formation with no monomer observable. SuhB monomer concentration was 16 nM.
(C) Rastergrams of the single-molecule traces with each row representing a single stalled TEC not containing the boxBAC element. Factor binding is represented as blue bars. Concentrations: 10 nM NusA-Cy5 or 50 nM NusG-Cy5.
(D) TEC-bound dwell-times of NusA and NusG were fitted to a single-exponential function using y=1-exp(-k*t), with coefficient k representing the off-rate. The errors represent 95 % confidence intervals of the fit. The number of fitted dwells (n) is indicated.
(E,F) Beeswarm plot with overlaid boxplot representing the longest binding event per trace (< 15 seconds shown) for each Cy5-labeled rrnTAC protein binding to the stalled TEC not containing the boxBAC element (E) or to the isolated boxBAC RNA (F), in the absence or presence of 400 nM of the rest of rrnTAC proteins, measured at 37 °C (except for the experiments with boxBAC RNA in presence of the rrnTAC proteins, which were measured at 30 °C). The number of molecules (n) and the number of events (shown in brackets) is indicated.
(G) Rastergrams of the single-molecule traces with each row representing a single boxBAC RNA molecule. Factor binding is represented as blue bars. Concentrations: 50 nM NusB-Cy5 or NusE-Cy5. Temperature: 21 °C. Salt: 20 mM KCl.
(H,I) Beeswarm plot with overlaid boxplot representing all protein-bound dwell times (H) or the number of binding events per trace (I) for the data represented in (G). The number of molecules (n) and the number of events (shown in brackets) is indicated.
(J) Fraction of boxBAC RNA molecules showing at least one protein binding event for the data represented in (G). The number of molecules (n) and the number of events (shown in brackets) is indicated.
(K) BoxBAC RNA-bound dwell times of NusE and NusB (same data as represented in panel (G)) were fitted to a single-exponential function using y=1-exp(-k*t), with coefficient k representing the off-rate. The errors represent 95 % confidence intervals of the fit. The number of fitted dwells (n) is indicated.

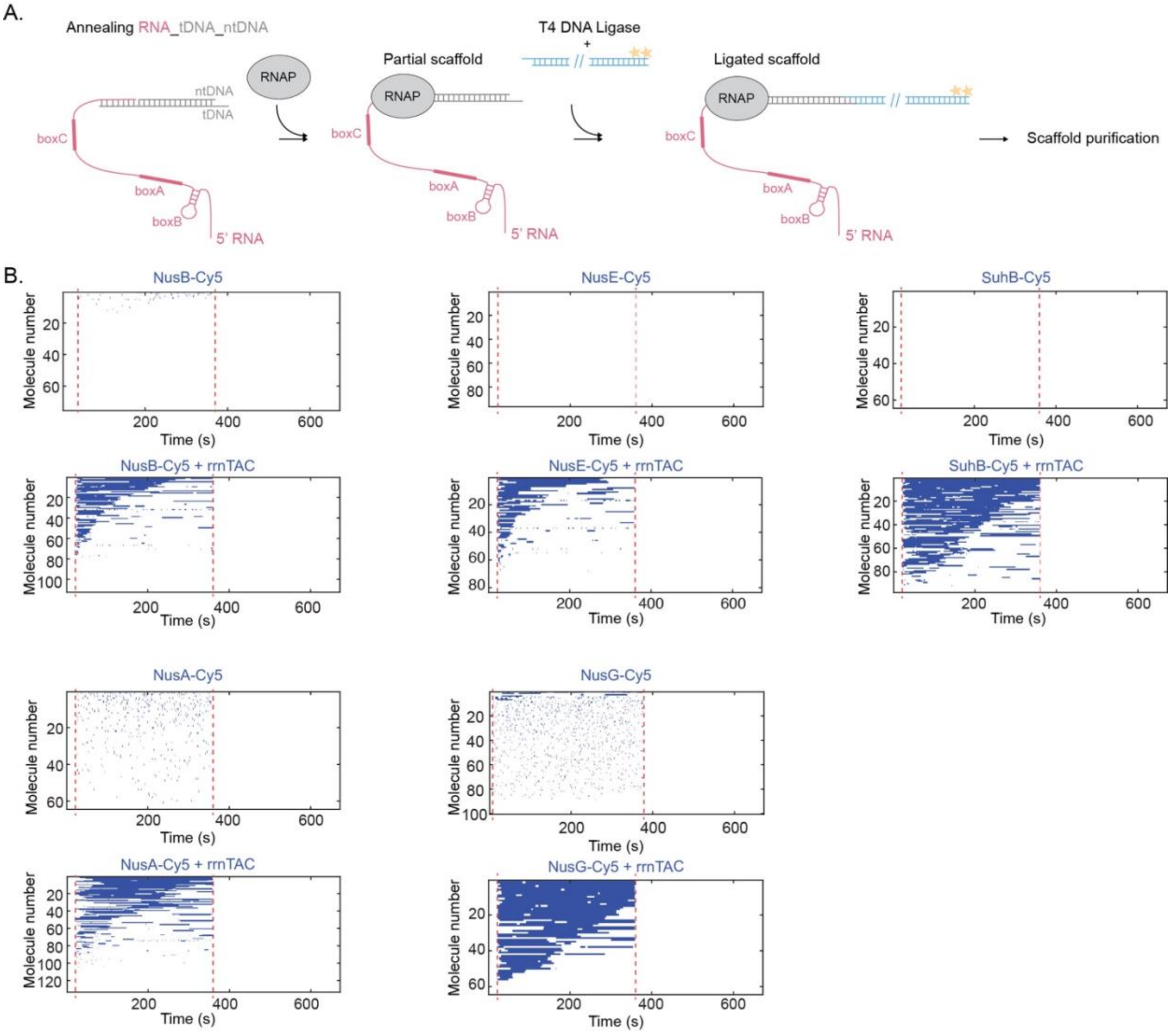


**Figure S3**. **Binding of rrnTAC proteins to the boxBAC-containing TEC.**

(A) Schematic representation of RNA:DNA:RNAP scaffold assembly pipeline.
(B) Rastergrams of the single-molecule traces with each row representing a single stalled TEC. Binding events of labeled proteins are shown as blue bars. Red dashed lines indicate reagents delivery: 1st injection – labeled rrnTAC protein in the absence or presence of 400 nM of the rest of rrnTAC proteins, 2nd injection – 200μM NTPs.

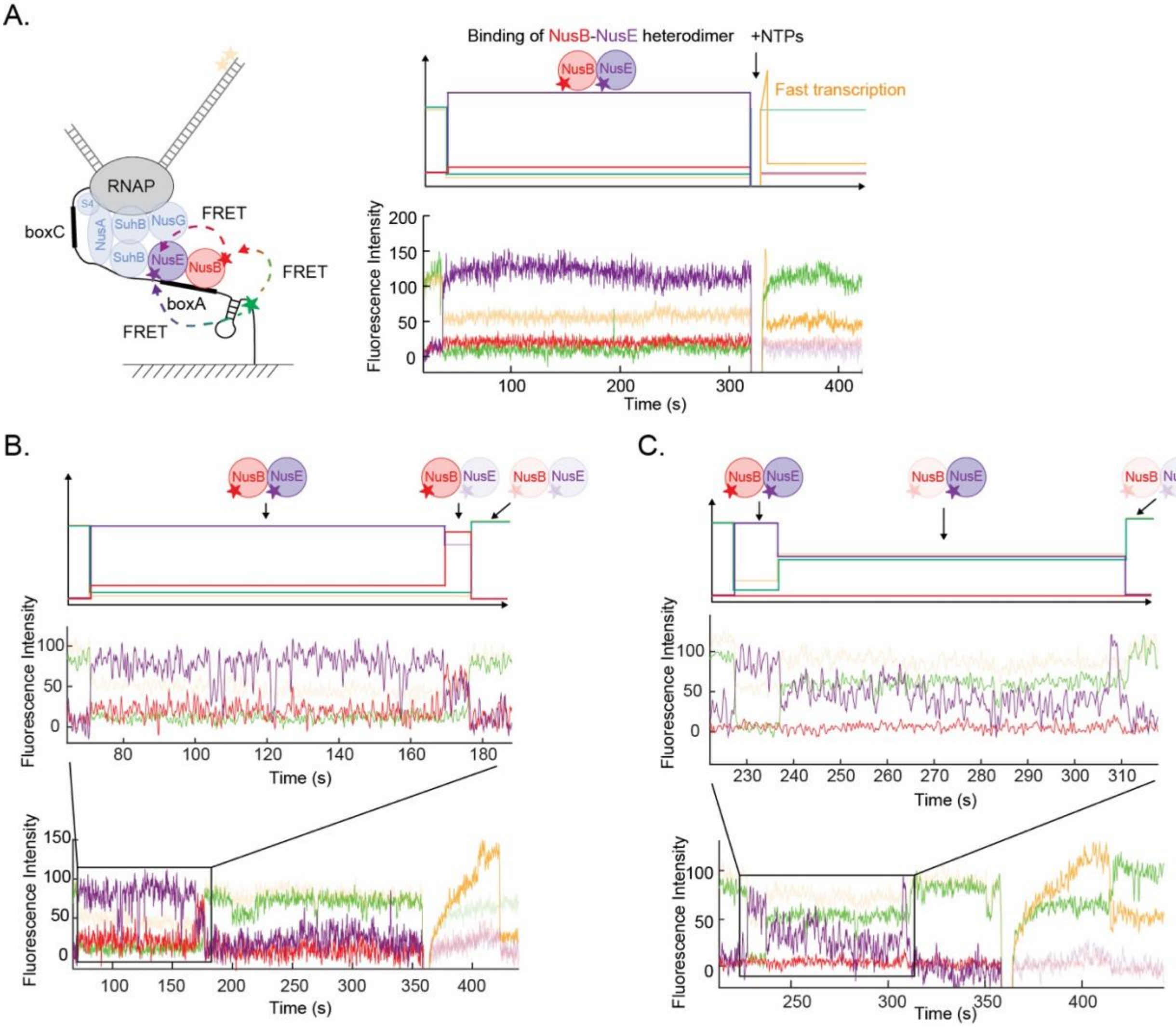


**Figure S4. NusB and NusE proteins bind boxBAC-containing TEC as a heterodimer.**

(A) Left panel: experimental setup to simultaneously track the binding dynamics of NusB-Cy5 and NusE-Cy5.5 with the rest of rrnTAC proteins unlabeled. Top right: schematic representation of a single-molecule trace showing binding of the NusB-Cy5/ NusE-Cy5.5 heterodimer. The co-recruitment of the two proteins at the same time shows a stable FRET event with the transfer from Cy3->Cy5->Cy5.5. Bottom right: representative single-molecule trace with binding of the NusB-Cy5.5/NusE-Cy5 heterodimer.
(B,C) Schematic (top) and experimental (bottom) single-molecule trace showing binding of the NusB-Cy5/ NusE-Cy5.5 heterodimer followed by NusE-Cy5.5 (B) or NusB-Cy5 (C) photobleaching.

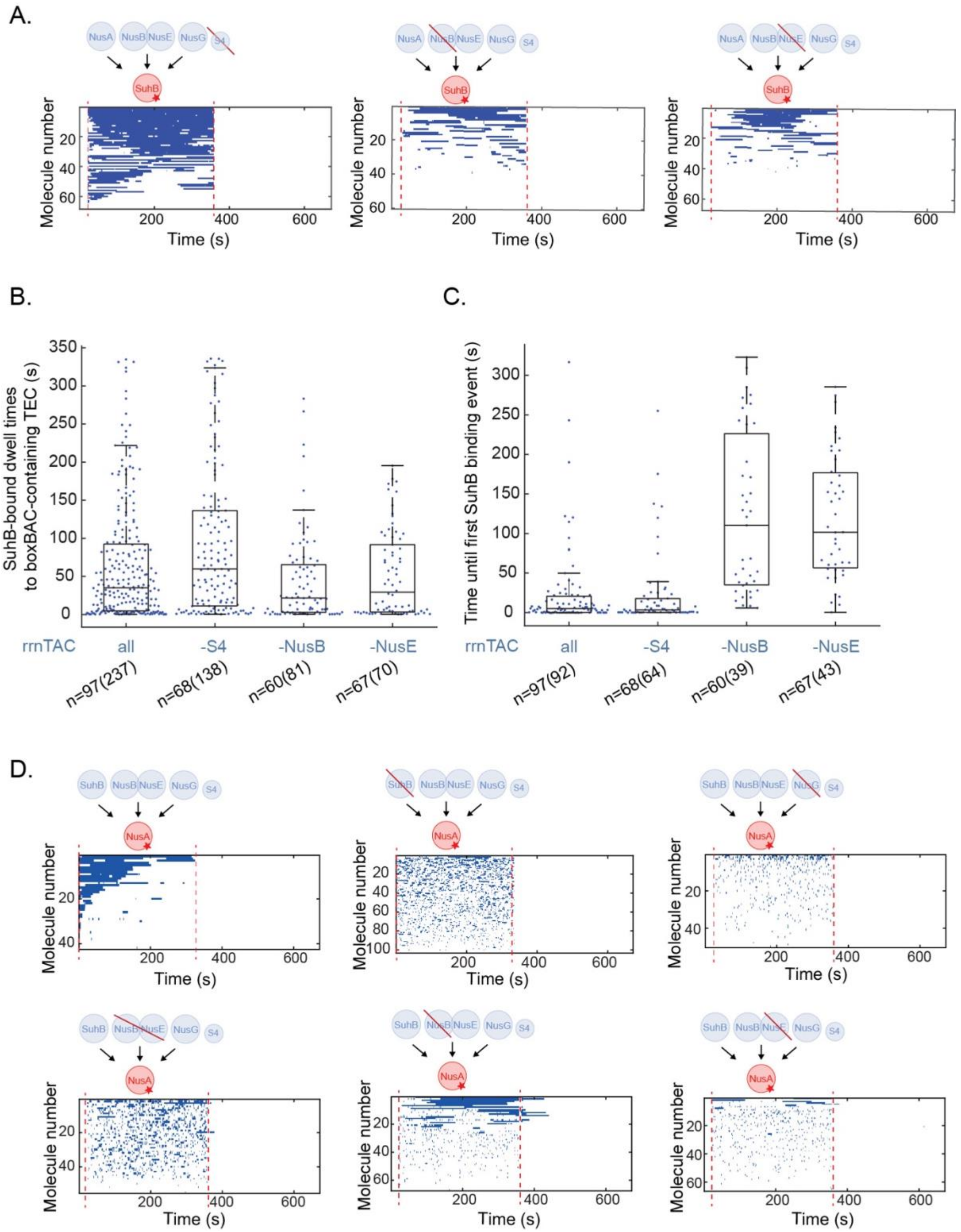


**Figure S5. All rrnTAC proteins (except S4) are required for efficient incorporation of SuhB and NusA.**

(A-C): S4 does not affect SuhB binding to rrnTAC and lacking NusB or NusE allows stable incorporation of SuhB into the boxBAC-containing TEC but with highly reduced recruitment rate. Rastergrams of the single-molecule traces with each row representing a single TEC (A). SuhB binding events are shown as blue bars; Beeswarm plot and overlaid boxplot of all SuhB-bound dwell times (B) or the time until the appearance of the first SuhB binding event (C) in presence of all rrnTAC proteins or omitting either S4, NusB or NusE. The number of molecules (n) and the number of events (shown in brackets) is indicated. (D) All rrnTAC proteins are required for fast and stable incorporation of NusA into the boxBAC-containing TEC. Rastergrams of the single-molecule traces with each row representing a single TEC. NusA binding events are shown as blue bars.

Red dashed lines indicate reagents delivery: 1$^{st}$ injection – labeled rrnTAC protein in the absence or presence of 400 nM of the subset of rrnTAC proteins, 2$^{nd}$ injection – 200µM NTPs.

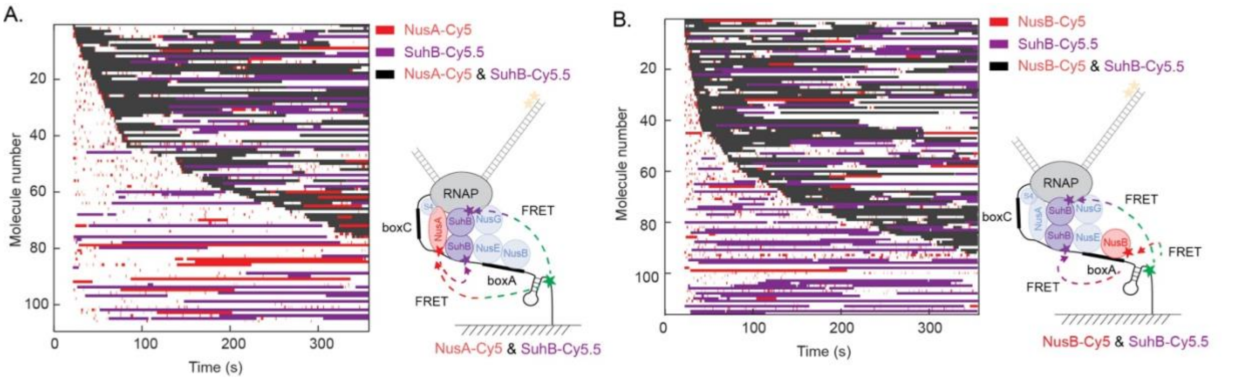


**Fig. S6. SuhB homodimer binds after NusA/NusB.**

(A,B) Rastergram of the single-molecule traces with each row representing a single stalled TEC. The observable binding events are presented as colored bars. For (A): NusA-Cy5 (red), SuhB-Cy5.5 (purple) and co-binding of both labeled proteins (black). Concentrations used: 10 nM NusA-Cy5, 50 nM SuhB-Cy5.5, 400 nM other unlabeled rrnTAC proteins. For (B): NusB-Cy5 (red), SuhB-Cy5.5 (purple) and co-binding of both labeled proteins (black). Concentrations used: 50 nM NusB-Cy5, 50 nM SuhB-Cy5.5, 400 nM other unlabeled rrnTAC proteins. Long binding events showing only SuhB-Cy5.5 binding (purple) or only NusA-Cy5/NusB-Cy5 binding (red), are also co-binding events but with the other protein being unlabeled due to incomplete labeling efficiency.

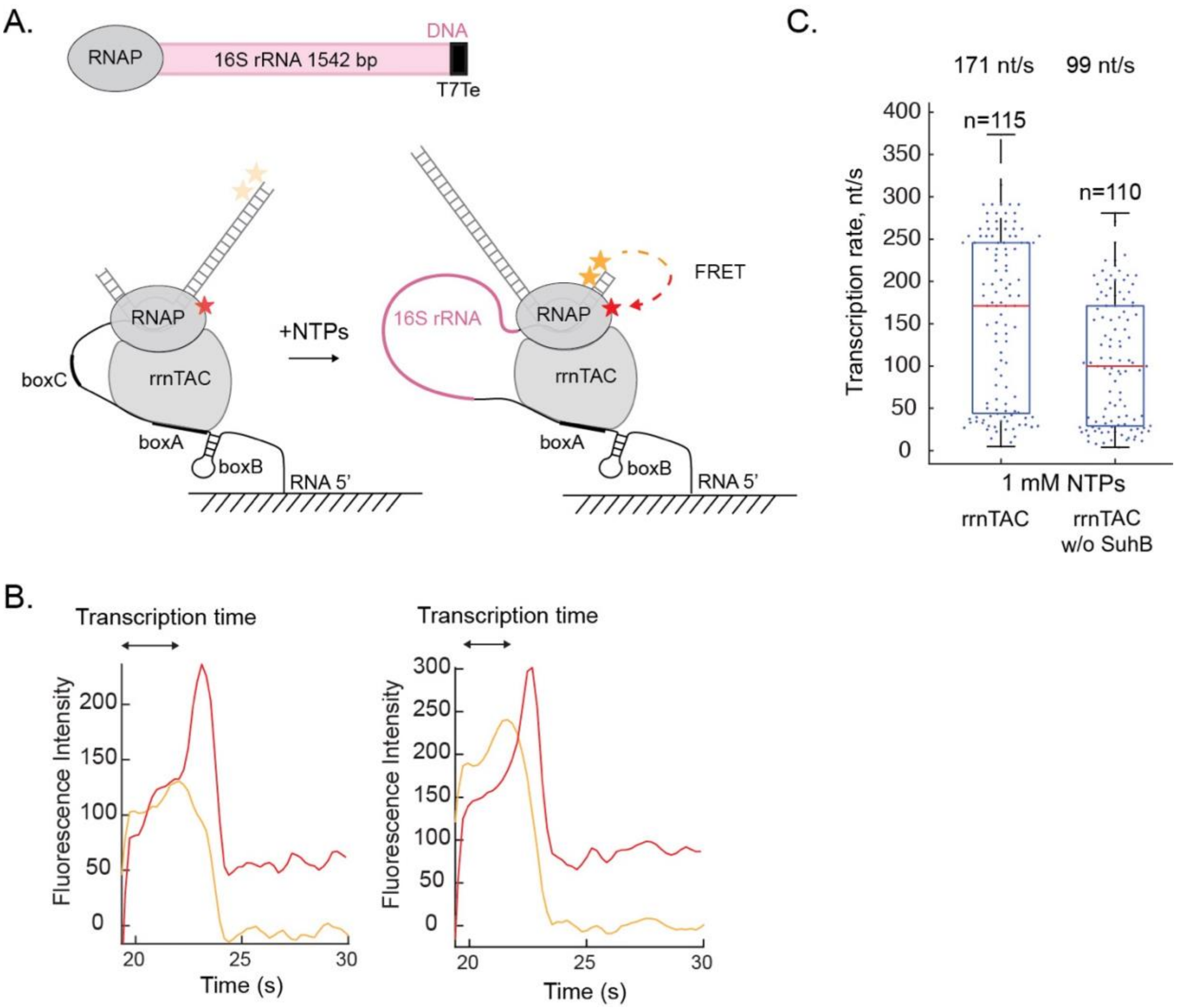


**Fig. S7. Stably assembled rrnTAC increases transcription speed about two-fold.**
(A) Upper panel: DNA template used for transcription. Bottom panel: experimental setup to study transcription speed with assembled rrnTAC. A FRET between Cy3.5 (DNA) and Cy5 (RNAP) indicates the end of transcription.
(B) Representative single-molecule traces showing Cy3.5 to Cy5 FRET. Measured transcription time is indicated.
(C) Beeswarm plot and overlaid boxplot of transcription speed of molecules with assembled rrnTAC, or with rrnTAC lacking SuhB at 37 °C. The number of molecules (n) is indicated.

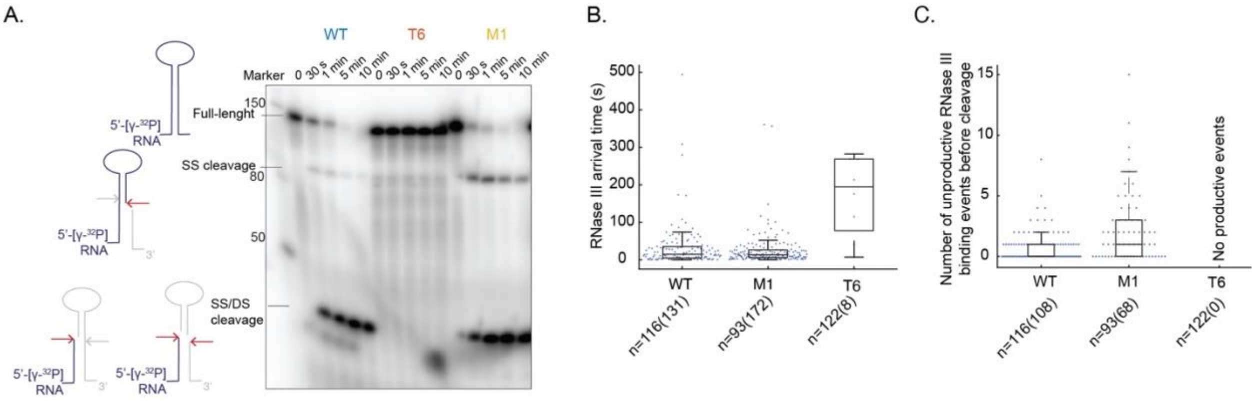


**Figure S8. RNase III cleaves native helix most efficiently.**

(A) Bulk RNase III cleavage assay with different RNA substrates (WT, M1, T6) at 3.5 mM $MgCl_2$, 37 °C, analyzed by denaturing gel electrophoresis.

(B-C) Single-molecule assay to quantify RNase III binding and cleavage of various mutant RNAs. Beeswarm plot and overlaid boxplot of RNase III arrival times (B), not including the arrival time to the first RNase III binding event and number of unproductive binding events before cleavage (C). Conditions: 20 nM RNase III-Cy5, 3.5 mM $MgCl_2$, 21 °C. The number of molecules (n) and the number of events (shown in brackets) is indicated.

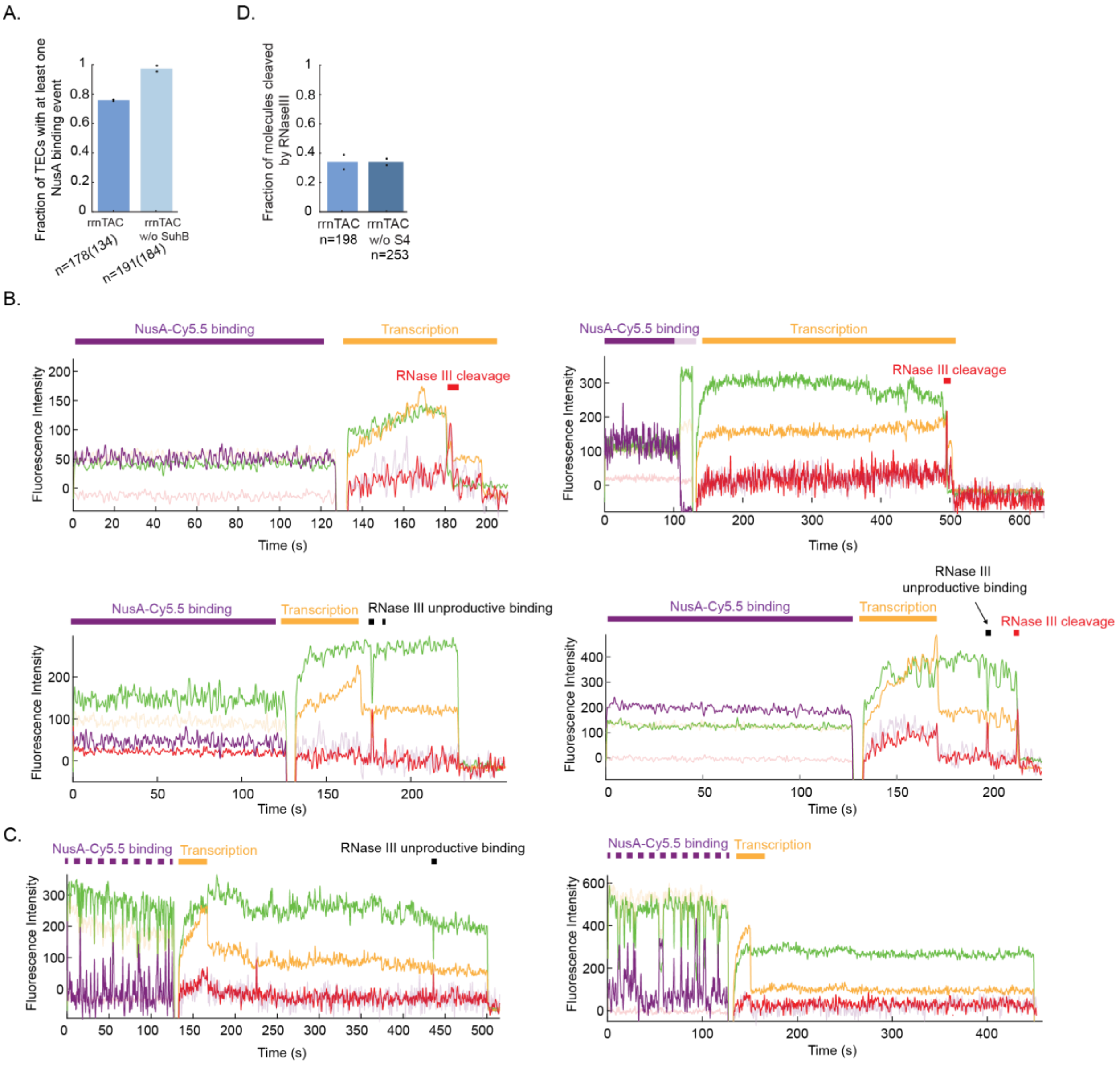


**Figure S9. rrnTAC-mediated co-transcriptional processing by RNase III**
(A) Fraction of TECs showing at least one NusA-Cy5.5 binding for assembled and not-assembled rrnTAC (lacking SuhB). Black dots represent values from two biological replicates.
(B) Additional representative single-molecule traces for the experiment shown in Fig. 6C (scenario in blue dashed box) for NusA-Cy5.5 binding in the presence of all rrnTAC proteins, demonstrating stable rrnTAC assembly, transcription and RNase III cleavage.
(C) Additional representative single-molecule traces for the experiment shown in Fig. 6C (scenario in pink dashed box) for NusA-Cy5.5 binding in the presence of all rrnTAC proteins but omitting SuhB, showing transient NusA-Cy5 binding.
(D) Fraction of nascent RNA molecules cleaved by RNase III. Black dots represent values from two biological replicates. Data in presence of rrnTAC is also shown in Fig. 6H.

The number of molecules (n) and the number of events (shown in brackets) is indicated.

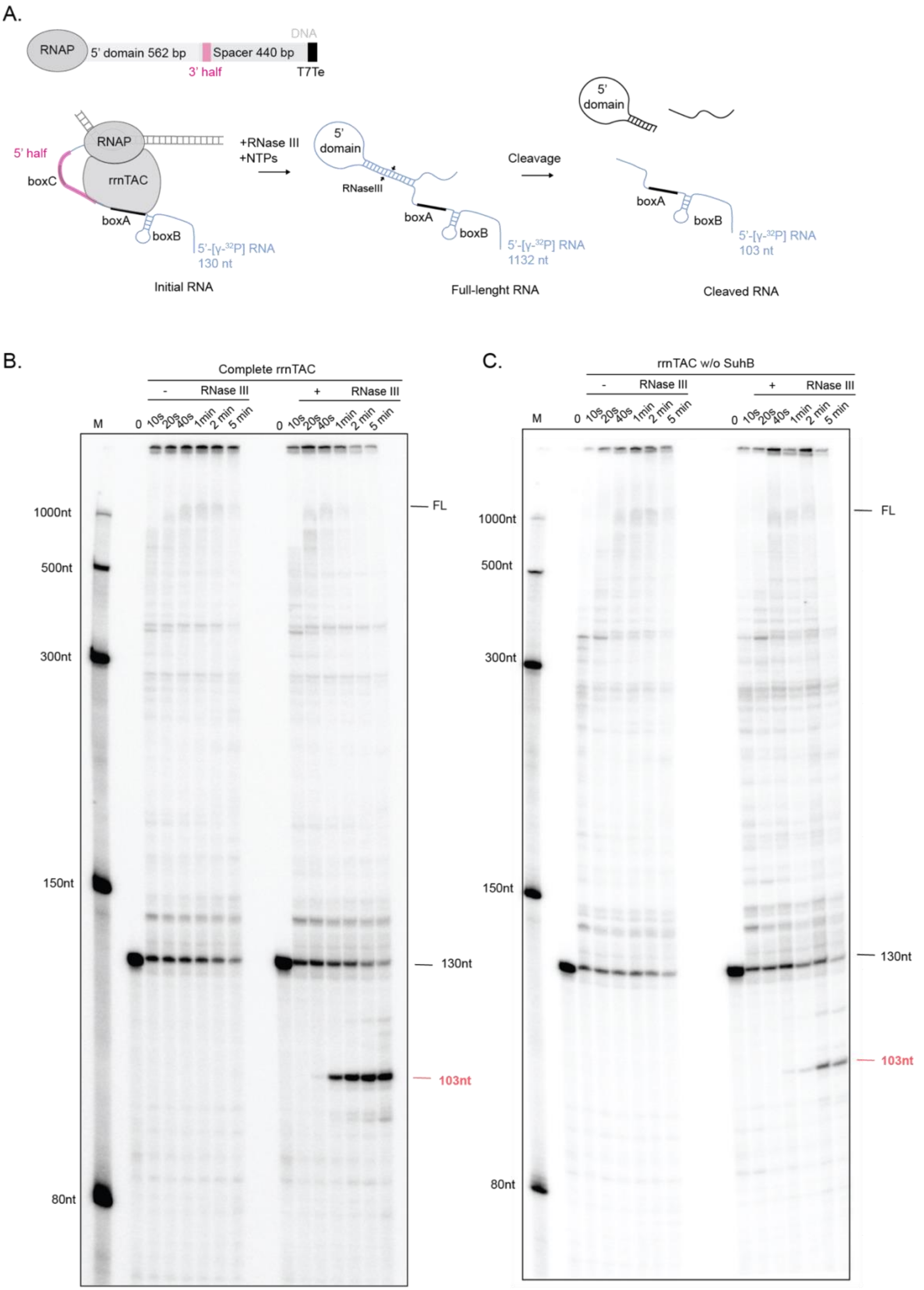


**Figure S10.** ***In vitro*** **single-round co-transcriptional RNase III cleavage assays**

(A) Upper panel: DNA template used for transcription encoding the 5’ domain of the 16S rRNA (562 bp), the 16S_23S spacer (440 bp) encoding the 3’ half of the RNase III substrate helix, and a T7 terminator. Bottom panel: schematic representation of the co-transcriptional RNase III processing assay. Radioactively labeled 5’-[$\gamma$-$^{32}$P] RNA products at different stages of the reaction are shown in blue.

(B), (C) Single-round co-transcriptional RNase III processing assays show that RNase III cleavage is specific and its efficiency increased in presence of all rrnTAC components. Radioactively labeled 5’-[$\gamma$-$^{32}$P] RNA was used to assemble an RNA:DNA scaffold bound by RNAP. Assembled complexes were purified with streptavidin beads prior to the assay to remove free 5’-[$\gamma$-$^{32}$P] RNA. For the reactions with the assembled rrnTAC, the purified scaffold was incubated for 10 minutes with 1 $\mu$M rrnTAC proteins at 37 ºC. Transcription was initiated by addition of 0.2 mM NTPs +/- 50 nM RNase III at 10 mM $MgCl_2$. Initial RNA (130 nt) and the full-length product are shown in black (1132 nt); the 5’ cleavage product by RNase III (103 nt) is shown in red.

**Supplementary Table 1. Overview of RNAs used in this study**

*DNA template for sequences 1-4:* T7promoter (18 nt) - G - revCompl_p109p44 (AC-rich, 25 nt) - revCompl_F2 (22 nt) - box BAC (61nt) - 20 nts native sequence - VS ribozyme minimal stem loop (24 nt)

*DNA template for sequence 7:* T7promoter (18 nt) - G - revCompl_F2 (22 nt) - box BAC (61nt) - CACTC - revCompl_p109p44 (AC-rich, 25 nt)

*DNA template for sequence 8-10:* T7promoter (18 nt) - G – revCompl_p0030 (30 nt) - A - long-range helix 5' half (31 nt)* - AAAAA - revCompl_F2 (22 nt) – AAAA - long-range helix 3' half (31 nt) - A - revCompl_p109p44 (AC-rich, 25 nt)

| # | ID | Sequence | Note |
|---|---|---|---|
| 1 | rnaAC001_not_VS_cleaved | ggacuaccaccacccaaccaacacaccaaccacuccaauuacauacaccgccgcugagaaaaagcgaagcggcacugcucuuuaacaauuuaucagacaaucugugugggcacucgaagauacggauucuaagggcgucgucgccccgaggauu | - |
| 2 | rnaAC001 | ggacuaccaccacccaaccaacacaccaaccacuccaauuacauacaccgccgcugagaaaaagcgaagcggcacugcucuuuaacaauuuaucagacaaucugugugggcacucgaagauacggauucu | Used for scaffold assembly |
| 3 | rnaAC011_not_VS_cleaved | ggacuaccaccacccaaccaacacaccaaccacuccaauuacauacaccgccgcugagaaaaagcgaagcggcacugcucuuuaacaauuuaucagacaaucuguguggguaagggcgucgucgccccgaggauu | - |
| 4 | rnaAC011 | ggacuaccaccacccaaccaacacaccaaccacuccaauuacauacaccgccgcugagaaaaagcgaagcggcacugcucuuuaacaauuuaucagacaaucugugugggg | Used for splint ligation |
| 5 | rnaAC012_Cy3 | cacucgaaga/Cy3/uacggauucuuaacguc | Purchased (IDT) |
| 6 | rnaAC013 | ggacuaccaccacccaaccaacacaccaaccacuccaauuacauacaccgccgcugagaaaaagcgaagcggcacugcucuuuaacaauuuaucagacaaucugugugggcacucgaaga/Cy3/uacggauucuuaacguc | Splint ligated: rnaAC011 and rnaAC012_Cy3; used for scaffold assembly |
| 7 | rnaAC002 | ggaaccacuccaauuacauacaccgccgcugagaaaaagcgaagcggcacugcucuuuaacaauuuaucagacaaucugugugggcacucacuaccaccacccaaccaacacacc | - |
| 8 | rnaAC015 | ggucacgaaagcugaguagucacgagucuucuauuuaucagacaaucugugugggcacucgaagaaaaaaaccacuccaauuacauacaccaaaaacuuuguagugcucacacagauugucugauagaaacuaccaccacccaaccaacacacc | WT |
| 9 | rnaAC016 | ggucacgaaagcugaguagucacgagucuucuauuuaucagacaa**A**cugugugggcacucgaagaaaaaaaccacuccaauuacauacaccaaaaacuuuguagugcucacacag**U**uugucugauagaaacuaccaccacccaaccaacacacc | M1 mutant (mutated nucleotides are shown in bold red) |
| 10 | rnaAC017 | ggucacgaaagcugaguagucacgagucuucuauuuaucagacaaucugugugggcacucgaagaaaaaaaccacuccaauuacauacaccaaaaacuuuguagug**GC**ca**GTTC**gauugucugauagaaacuaccaccacccaaccaacacacc | T6 mutant (Warner et al., 2023) (mutated nucleotides are shown in bold red) |

**Supplementary Table 2. Overview of linear DNA fragments used for scaffold assembly or for formation of stalled TEC**

*Partial scaffolds:*

| | ID | Sequence 5’ to 3’ |
|---|---|---|
| Partial scaffold 1 | tDNA-prAC154 | 5’Pi ctcttgagaccttggtatttatttttttgttaagataagggatagggaagaatccgt |
| | ntDNA-prAC155 | ccctatcccttatcttaacaaaaaaataaataccaaggtctca |
| Partial scaffold 2 | tDNA-prAC199 | 5'Pi ctcttgagaccttggtattcatttttcgtcttgcgacgttaag |
| | ntDNA-prAC198 | caagacgaaaaatgaataccaaggtctca |

*Connectors:*

native – BsaI – native – 5’ domain 16S rRNA / full 16S rRNA – native – RNase III substrate helix (3’ half) – native 16S_23S rRNA spacer – T7Te – ssDNA (complementary to p0088_2xCy3.5, to prAC143_biotin*)
*XhoI restriction site generated with hybridization of prAC143 to connector DNA is underlined

| | ID | Sequence 5’ to 3’ |
|---|---|---|
| 5’domain_spacer_T7Te | prAC217 | cttttccttctgttggtctcaagagtgaacacgtaattcattacgaagtttaattctttgagcgtcaaactttttaaattgaagagtttgatcatggctcagattgaacgctggcggcaggcctaacacatgcaagtcgaacggtaacaggaagaagcttgcttctttgctgacgagtggcggacgggtgagtaatgtctgggaaactgcctgatggagggggataactactggaaacggtagctaataccgcataacgtcgcaagaccaaagagggggaccttcgggcctcttgccatcggatgtgcccagatgggattagctagtaggtggggtaacggctcacctaggcgacgatccctagctggtctgagaggatgaccagccacactggaactgagacacggtccagactcctacgggaggcagcagtggggaatattgcacaatgggcgcaagcctgatgcagccatgccgcgtgtatgaagaaggccttcgggttgtaaagtactttcagcggggaggaagggagtaaagttaatacctttgctcattgacgttacccgcagaagaagcaccggctaactccgtgccagcagccgcggtaatacggagggtgcaagcgttaatcccttaaagaagcgtactttgtagtgctcacacagattgtctgatagaaagtgaaaagcaaggcgtttacgcgttgggagtgaggctgaagagaataaggccgttcgctttctattaatgaaagctcaccctacacgaaaatatcacgcaacgcgtgataagcaattttcgtgtccccttcgtctagaggcccaggacaccgccctttcacggcggtaacaggggttcgaatcccctaggggacgccacttgctggtttgtgagtgaaagtcgccgaccttaatatctcaaaactcatcttcgggtgatgtttgagatttttgctctttaaaatctggatcaagctgaaaattgaaacactgaacaacgagagttgttcgtgagtctctcaaattttcgcaacacgatgatgaatcgaaagaaacatcttcgggttgtgaTAATCACACTGGCTCACCTTCGGGTGGGCCTTTCTGCGTTTATGGGATGGTAATTTGGTGAGTATGATTAAGGatctc<u>CTCGAG</u>gactagctg |
| 16SrRNA_spacer_T7Te | prAC185 | cttttccttctgttggtctcaagagtgaacacgtaattcattacgaagtttaattctttgagcgtcaaactttttaaattgaagagtttgatcatggctcagattgaacgctggcggcaggcctaacacatgcaagtcgaacggtaacaggaagaagcttgcttctttgctgacgagtggcggacgggtgagtaatgtctgggaaactgcctgatggagggggataactactggaaacggtagctaataccgcataacgtcgcaagaccaaagagggggaccttcgggcctcttgccatcggatgtgcccagatgggattagctagtaggtggggtaacggctcacctaggcgacgatccctagctggtctgagaggatgaccagccacactggaactgagacacggtccagactcctacgggaggcagcagtggggaatattgcacaatgggcgcaagcctgatgcagccatgccgcgt |

| | | |
|---|---|---|
| | | gtatgaagaaggccttcgggttgtaaagtactttcagcggggaggaagggagtaaagtt<br>aatacctttgctcattgacgttacccgcagaagaagcaccggctaactccgtgccagcag<br>ccgcggtaatacggagggtgcaagcgttaatcggaattactgggcgtaaagcgcacgc<br>aggcggtttgttaagtcagatgtgaaatccccgggctcaacctgggaactgcatctgatac<br>tggcaagcttgagtctcgtagaggggggtagaattccaggtgtagcggtgaaatgcgtag<br>agatctggaggaataccggtggcgaaggcggccccctggacgaagactgacgctcag<br>gtgcgaaagcgtggggagcaaacaggattagataccctggtagtccacgccgtaaacg<br>atgtcgacttggaggttgtgcccttgaggcgtggcttccggagctaacgcgttaagtcgac<br>cgcctggggagtacggccgcaaggttaaaactcaaatgaattgacgggggcccgcac<br>aagcggtggagcatgtggtttaattcgatgcaacgcgaagaaccttacctggtcttgacat<br>ccacggaagttttcagagatgagaatgtgccttcgggaaccgtgagacaggtgctgcatg<br>gctgtcgtcagctcgtgttgtgaaatgttgggttaagtcccgcaacgagcgcaacccttatc<br>ctttgttgccagcggtccggccgggaactcaaaggagactgccagtgataaactggagg<br>aaggtggggatgacgtcaagtcatcatggcccttacgaccagggctacacacgtgctac<br>aatggcgcatacaaagagaagcgacctcgcgagagcaagcggacctcataaagtgc<br>gtcgtagtccggattggagtctgcaactcgactccatgaagtcggaatcgctagtaatcgt<br>ggatcagaatgccacggtgaatacgttcccgggccttgtacacaccgcccgtcacacca<br>tgggagtgggttgcaaaagaagtaggtagcttaaccttcgggagggcgcttaccactttgt<br>gattcatgactggggtgaagtcgtaacaaggtaaccgtaggggaacctgcggttggatc<br>acctccttaccttaaagaagcgtactttgtagtgctcacacagattgtctgatagaaagtga<br>aaagcaaggcgtttacgcgttgggagtgaggctgaagagaataaggccgttcgctttcta<br>ttaatgaaagctcaccctacacgaaaatatcacgcaacgcgtgataagcaattttcgtgtc<br>cccttcgtctagaggcccaggacaccgccctttcacggcggtaacaggggttcgaatccc<br>ctaggggacgccacttgctggtttgtgagtgaaagtcgccgaccttaatatctcaaaactc<br>atcttcgggtgatgtttgagatttttgctctttaaaaatctggatcaagctgaaaattgaaaca<br>ctgaacaacgagagttgttcgtgagtctctcaaattttcgcaacacgatgatgaatcgaaa<br>gaaacatcttcgggttgtgaTAATCACACTGGCTCACCTTCGGGTGGG<br>CCTTTCTGCGTTTATGGGATGGTAATTTGGTGAGTATGATTAA<br>GGatctcCTCGAGgactagctg |
| 16SrRNA _T7Te | prAC234 | cttttccttctgttggtctcaagagtgaacacgtaattcattacgaagtttaattctttgagcgtc<br>aaacttttaaattgaagagtttgatcatggctcagattgaacgctggcggcaggcctaaca<br>catgcaagtcgaacggtaacaggaagaagcttgcttctttgctgacgagtggcggacgg<br>gtgagtaatgtctgggaaactgcctgatggagggggataactactggaaacggtagcta<br>ataccgcataacgtcgcaagaccaaagagggggaccttcgggcctcttgccatcggatg<br>tgcccagatgggattagctagtaggtggggtaacggctcacctaggcgacgatccctag<br>ctggtctgagaggatgaccagccacactggaactgagacacggtccagactcctacgg<br>gaggcagcagtggggaatattgcacaatgggcgcaagcctgatgcagccatgccgcgt<br>gtatgaagaaggccttcgggttgtaaagtactttcagcggggaggaagggagtaaagtt<br>aatacctttgctcattgacgttacccgcagaagaagcaccggctaactccgtgccagcag<br>ccgcggtaatacggagggtgcaagcgttaatcggaattactgggcgtaaagcgcacgc<br>aggcggtttgttaagtcagatgtgaaatccccgggctcaacctgggaactgcatctgatac<br>tggcaagcttgagtctcgtagaggggggtagaattccaggtgtagcggtgaaatgcgtag<br>agatctggaggaataccggtggcgaaggcggccccctggacgaagactgacgctcag<br>gtgcgaaagcgtggggagcaaacaggattagataccctggtagtccacgccgtaaacg<br>atgtcgacttggaggttgtgcccttgaggcgtggcttccggagctaacgcgttaagtcgac<br>cgcctggggagtacggccgcaaggttaaaactcaaatgaattgacgggggcccgcac<br>aagcggtggagcatgtggtttaattcgatgcaacgcgaagaaccttacctggtcttgacat<br>ccacggaagttttcagagatgagaatgtgccttcgggaaccgtgagacaggtgctgcatg<br>gctgtcgtcagctcgtgttgtgaaatgttgggttaagtcccgcaacgagcgcaacccttatc<br>ctttgttgccagcggtccggccgggaactcaaaggagactgccagtgataaactggagg<br>aaggtggggatgacgtcaagtcatcatggcccttacgaccagggctacacacgtgctac<br>aatggcgcatacaaagagaagcgacctcgcgagagcaagcggacctcataaagtgc<br>gtcgtagtccggattggagtctgcaactcgactccatgaagtcggaatcgctagtaatcgt<br>ggatcagaatgccacggtgaatacgttcccgggccttgtacacaccgcccgtcacacca<br>tgggagtgggttgcaaaagaagtaggtagcttaaccttcgggagggcgcttaccactttgt<br>gattcatgactggggtgaagtcgtaacaaggtaaccgtaggggaacctgcggttggatc |

| | | |
|---|---|---|
| | | acctccttaTAATCACACTGGCTCACCTTCGGGTGGGCCTTTCTGCGTTTATGGGATGGTAATTTGGTGAGTATGATTAAGGatctcCTCGAGgactagctg |

*DNA fragments used for formation of stalled TEC*

ssDNA - upstream sequence – P1promoter – native sequence – boxBAC – native sequence – 5'domain – T7Te

| ID | Sequence 5' to 3' |
|---|---|
| prAC235 | TCACGAAAGCTGAGTAGTCACGAGTCTTCTctggcagttttaggctgatttggttgaatgttgcgcggtcagaaaattattttaaatttcctcttgtcaggccggaataactccctataatgcgccaccactgacacggaacaacggcaaacacgccgccgggtcagcggggttctcctgagaactccggcagagaaagcaaaaataaatgcttgactctgtagcgggaaggcgtattatgcacaccccgcgccgctgagaaaaagcgaagcggcactgctcttttaacaatttatcagacaatctgtgtgggcactcgaagatacggattcttaacgtcgcaagacgaaaaatgaataccaagtctcaagagtgaacacgtaattcattacgaagtttaattctttgagcgtcaaacttttaaattgaagagtttgatcatggctcagattgaacgctggcggcaggcctaacacatgcaagtcgaacggtaacaggaagaagcttgcttctttgctgacgagtggcggacgggtgagtaatgtctgggaaactgcctgatggagggggataactactggaaacggtagctaataccgcataacgtcgcaagaccaaagagggggaccttcgggcctcttgccatcggatgtgcccagatgggattagctagtaggtggggtaacggctcacctaggcgacgatccctagctggtctgagaggatgaccagccacactggaactgagacacggtccagactcctacgggaggcagcagtggggaatattgcacaatgggcgcaagcctgatgcagccatgccgcgtgtatgaagaaggccttcgggttgtaaagtactttcagcggggaggaagggagtaaagttaatacctttgctcattgacgttacccgcagaagaagcaccggctaactccgtgccagcagccgcggtaatacggagggtgcaagcgttaatcTAATCACACTGGCTCACCTTCGGGTGGGCCTTTCTGCGTTTAT |

ssDNA – upstream sequence – P1promoter – revCompl_p109p44 (AC-rich, 25nts) - revCompl_F2 (22 nt)– native sequence 3' domain 16S rRNA – T7Te – ssDNA

| ID | Sequence 5' to 3' |
|---|---|
| T13 | TCACGAAAGCTGAGTAGTCACGAGTCTTCTctggcagttttaggctgatttggttgaatgttgcgcggtcagaaaattattttaaatttcctcttgtcaggccggaataactccctataatgcgccaccactaccaccacccaaccaacacaccaaccactccaattacatacacctgacgggggcccgcacaagcggtggagcatgtgtcaccacgtgctacaatggcgcatacaaagagaagcgacctcgcgagagcaagcggacctcataaagtgcgtcgtagtccggattggagtctgcaactcgactccatgaagtcggaatcgctagtaatcgtggatcagaatgccacggtgaatacgttcccgggccttgtacaccctatcccttatcttaacggctccttttggagccttttttttggagattttctaaaacgaaaggctcagtcgaaagactgggcctttcgttttatctTAATCACACTGGCTCACCTTCGGGTGGGCCTTTCTGCGTTTATgggatggtaatttggtgagtatgattaagg |

**Supplementary Table 3. Overview of oligonucleotides used in this study**

| **Oligonucleotide** | **DNA sequence from 5' to 3' end** |
|---|---|
| p0088-2xCy3.5 | 5'Cy3.5-GGGATGGTAATTTGGdT(Cy3.5)GAGTATGATTAAGG |
| prAC143_biotin | ATCTCCTCGAGGACTAGCTG-3' Biotin |
| p0141 | GGTGTGTTGGTTGGGTGGTGGTAGTTGTGGAATTGTGAGCGGATA A |
| p0109-biotin | 5'BiotinTEG-TTATCCGCTCACAATTCCACA |
| p0155 | TGTGGAATTGTGAGCGGATAAGGTGTGTTGGTTGGGTGGTGGTAGT |
| p0154-biotin | TTATCCGCTCACAATTCCACA-3'BiotinTEG |
| p066-Cy3 | 5'Cy3-GGTGTATGTAATTGGAGTGGTT |
| p0150_Cy3p5 | 5'Cy3.5-AGAAGACTCGTGACTACTCAGCTTTCGTGA |
| prAC197 | GACGTTAAGAATCCGTATCTTCGAGTGCCCACACAGATTGTCTGATAAATTGT TAAAGAG |

**Supplementary Material:**

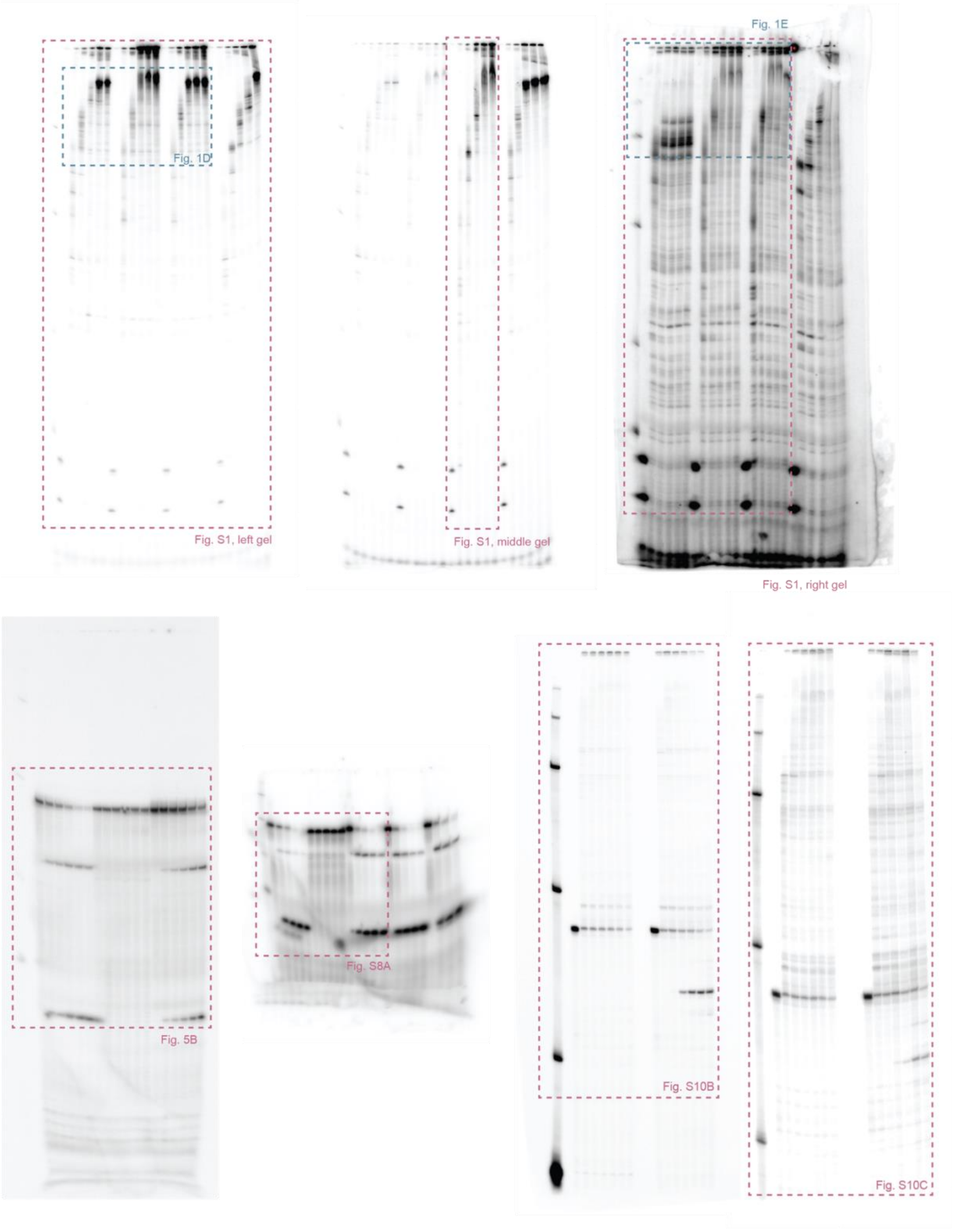


**Uncropped gels from Fig. 1, 5, S1, S8, S10**
